# Tissue-level division of labor is coordinated across organs through universal programs and context-specific adaptations

**DOI:** 10.1101/2025.09.16.676484

**Authors:** Shira Yonassi, Noa Goldenberg, Avia Walfisch, Eitan Cohen, Einat Shaer Tamar, Miri Adler

## Abstract

Fibroblasts, macrophages, and endothelial cells are common to most mammalian tissues, where they support homeostasis, repair, and immune regulation. Yet how they coordinate their functions across diverse tissue environments remains unclear. Here, we analyze single-cell RNA sequencing data from 14 human tissues using Pareto optimality framework and identify 16 archetypes - specialized transcriptional programs representing functional tradeoffs. These include universal archetypes shared across tissues and tissue-specific archetypes shaped by local context. We find that tissues align along a continuous axis of archetype distributions, from metabolically active to barrier organs, reflecting coordinated functional adaptation. Using ligand-receptor enrichment mapping, we reveal task-specific crosstalk across cell types, suggesting that intercellular communication underlies supportive division of labor. Applying this analysis to mouse tissues, we find similar patterns, indicating evolutionary conserved tradeoffs. These findings provide a framework for understanding how division of labor is coordinated in health and how its dysregulation may contribute to disease.

## Introduction

Our tissues depend on the coordinated activity of diverse cell populations to maintain physiological function. Typically, each tissue is organized around one or more primary cell types that perform the tissue’s defining function, including neurons in the brain, hepatocytes in the liver, and cardiomyocytes in the heart. These primary cells are supported by other cell types that provide structural, metabolic, and immunological support^1–3^. Among the most ubiquitous and indispensable of these are fibroblasts, macrophages, and capillary endothelial cells. These cell types are present in nearly every tissue where they contribute to essential tissue-level functions. Fibroblasts maintain extracellular structure and regulate wound healing^4^, macrophages regulate immune surveillance and inflammation^5,6^, and endothelial cells control nutrient and oxygen delivery^7,8^. Together with the tissue’s primary cells, these supportive cells form a minimal unit of tissue functionality^1,3,9^ (Fig. 1A-B). Their essentiality is further demonstrated by their key roles in diseases characterized by disrupted homeostasis, including cancer^10–14^, fibrosis^15–19^, and chronic inflammation^20–22^.

**Figure 1:**
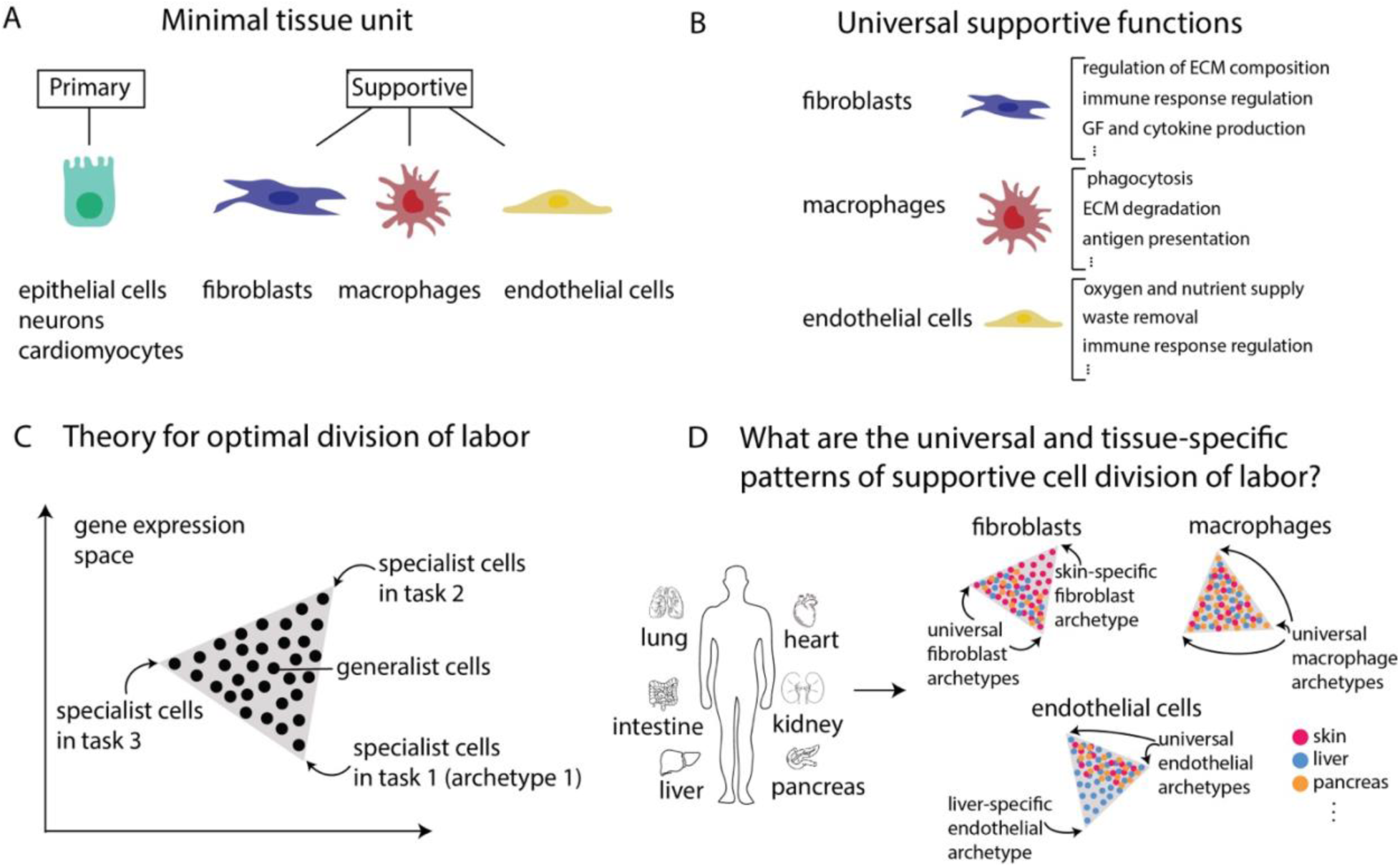
Mapping universal and tissue-specific patterns of division of labor among supportive cell populations. (A) Schematics of a minimal tissue unit composed of a primary cell type and supportive cells that are common to most tissues: fibroblasts, macrophages, and endothelial cells. (B) Supportive cell types perform multiple functions to facilitate the primary tissue function. (C) Based on evolutionary tradeoff theory, individual cells are predicted to fill in a polytope in gene expression space (triangle, tetrahedron, etc.). Specialist cells are found near the vertices (archetypes) and generalist cells are in the middle of the polytope. (D) Universal and tissue-specific archetypes can be defined by comparing distributions of common cell populations within the polytope across tissues.

Each supportive population is required to perform a wide range of functions, many of which are incompatible at the single-cell level (Fig. 1B). For example, a fibroblast may not efficiently perform both matrix deposition and immune signaling at full capacity simultaneously^23–26^. To resolve such constraints, cells often divide labor, resulting in functional heterogeneity within a differentiated cell type. Division of labor has been demonstrated in primary cell types of tissues such as the liver^27^ and intestine^28,29^, and has also been detected in fibroblasts, macrophages, and other cell types^25,30–33^. However, the principles and organization underlying these specializations remain poorly understood.

To interpret this heterogeneity, we previously developed a framework based on evolutionary tradeoffs, using Pareto optimality theory to uncover cell states that represent optimal solutions to conflicting functional demands^29^. This framework predicts that the gene expression profiles of cells performing multiple incompatible tasks are constrained within a polytope, whose vertices represent archetypes, specialist cells optimized for distinct functions, while interior points in the polytope represent generalists balancing multiple functions (Fig. 1C). We previously applied this framework to various cell populations, uncovering tradeoff-driven specializations and distinguishing between extrinsically imposed versus self-organized strategies that regulate this specialization^29,30^.

For common cell types such as fibroblasts, macrophages, and endothelial cells, a comprehensive framework is needed to explain whether their division of labor follows universal principles or is shaped by tissue-specific contexts. While these cells share core transcriptional programs, they also exhibit context-specific expression profiles influenced by environmental factors such as oxygen availability, turnover rates, and immune activity^34–37^. Previous studies have made substantial progress in identifying shared and tissue-specific subpopulations of common supportive cells by applying clustering approaches to large single-cell datasets. These studies revealed transcriptional heterogeneity and enabled the construction of pan-tissue atlases that describe conserved states across organs^2,38–47^. However, the biological interpretation of these clusters and the principles governing why these cells partition into distinct subpopulations often remains unclear. Clustering methods group cells by similarity, but do not directly explain why certain transcriptional profiles emerge or how they reflect underlying tradeoffs. Moreover, when gene expression varies continuously between multiple extremes, as in epithelial-mesenchymal plasticity^48^ or macrophage polarized states^49^, clustering may yield arbitrary partitions and mask gradual transitions.

Pareto optimality can address this by interpreting cellular variation in terms of functional tradeoffs, revealing archetypes as optimal specialists and allowing generalist states to emerge naturally as combinations of these extremes. This approach provides a mechanistic explanation for population structure and reveals the axes of functional tradeoffs that drive specialization. Moreover, it is especially beneficial in cases lacking clear-cut clusters, offering a principled, quantitative perspective into cellular heterogeneity.

Here, we apply and extend the Pareto optimality framework to single-cell RNA sequencing (scRNA-seq) data from 14 human tissues, focusing on fibroblasts, macrophages, and endothelial cells. We identify both universal archetypes, shared across tissues, and tissue-specific archetypes, shaped by local microenvironments (Fig. 1D). We uncover conserved axes of variation, revealing how supportive cells tailor their specialization according to tissue demands. We also explore ligand-receptor communication networks among archetypes, finding that supportive cells communicate through task-specific crosstalk, coordinating their functions in a process-dependent manner. Finally, we examine the conservation of these archetypes in mouse tissues, revealing evolutionary stability of core tradeoffs and specialization strategies. Our approach uncovers a unified framework for understanding how essential supportive cell populations organize, specialize, and coordinate functions across tissues.

## Results

### Common supportive cell populations exhibit multiple functional archetypes

To systematically investigate how supportive cell types divide labor across tissues, we analyzed single-cell RNA sequencing (scRNA-seq) data of macrophages, fibroblasts, and endothelial cells from 14 human tissues, including lung, liver, heart, intestines, and skin (data from^47^, Methods). We applied the Pareto Task Inference algorithm (ParTI^50^, Methods) to identify archetypes - extremal transcriptional states that represent cells optimized for distinct functional specializations. In this framework, archetypes correspond to the vertices of a polytope in gene expression space, with nearby cells interpreted as specialists and interior cells as generalists balancing multiple tasks (Fig. 1C). To define the major functional axes for each cell type, we pooled cells from all tissues and selected highly variable genes (Methods).

We find five archetypes of macrophage specialization (AM1-AM5, *p* < 0.001; Fig. 2A, Fig. S1A), each enriched for various gene programs. Gene enrichment analysis indicated that these archetypes reflect specialization in: phagocytosis and immune signaling; immune resolution and homeostatic maintenance; cytokine production and antigen presentation; pro-inflammatory and antimicrobial responses; and mitochondrial metabolism and protein synthesis. In fibroblasts, we similarly identified five archetypes (AF1-AF5, *p* < 0.001; Fig. 2B, Fig. S1B), associated with functions including: wound healing and extracellular matrix (ECM) organization; metabolic activity and regulation of cell identity and growth; ECM production, tissue remodeling, and contractility; immune signaling; and mitochondrial and biosynthetic processes. Endothelial cells exhibited six archetypes (AE1-AE6, *p* < 0.001; Fig. 2C, Fig. S1C), reflecting specialization in: cell migration and angiogenesis; barrier regulation and endocytosis; metabolic reprogramming and angiogenesis; mitochondrial energy production, permeability and innate immunity; adaptive immunity; and protein synthesis. Each archetype is defined by a set of 172-1415 enriched genes (listed in Tables S1-S3) representing additional functions.

**Figure 2:**
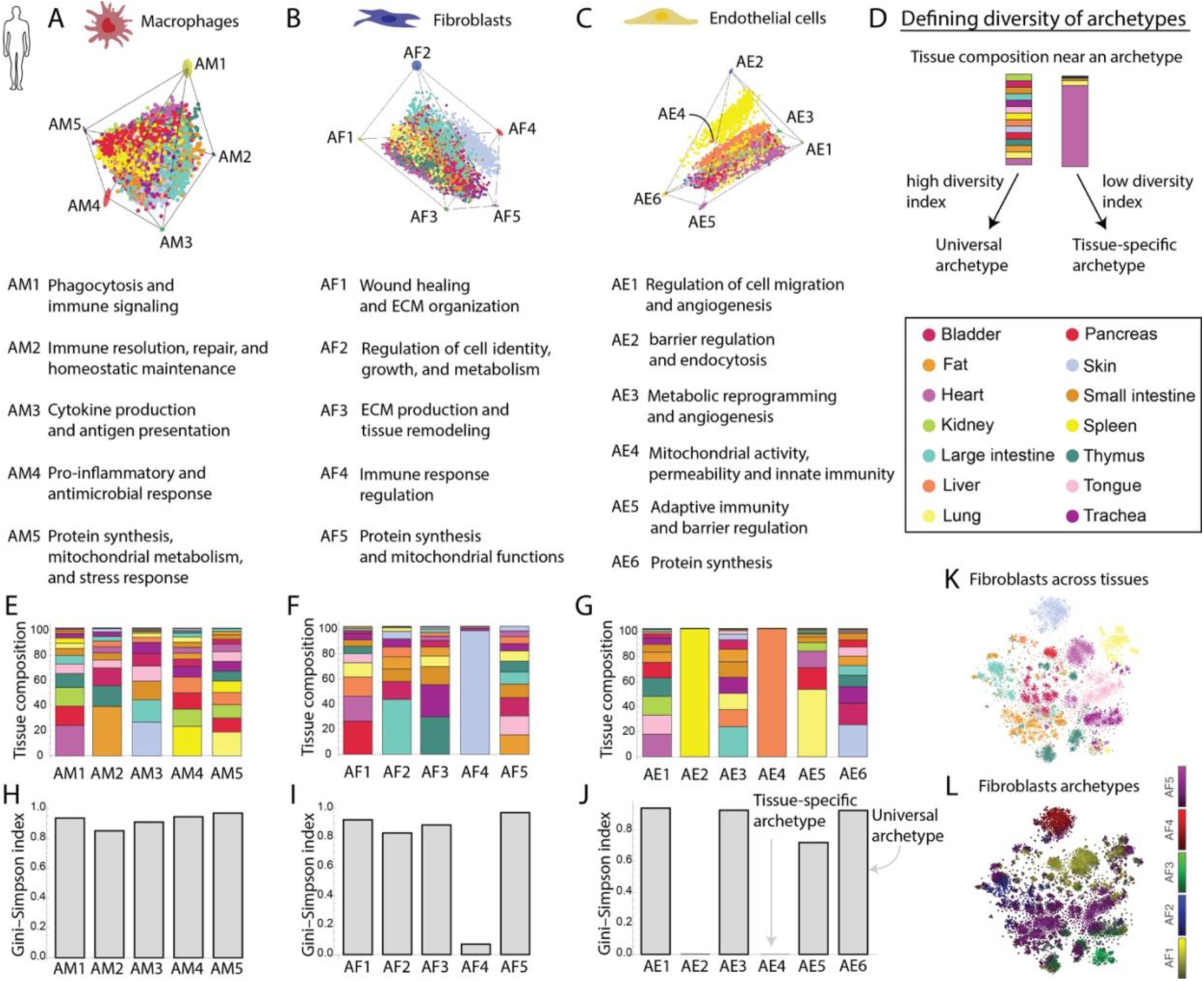
Identifying universal and tissue-specific archetypes in supportive populations using Pareto optimality. (A-C) Polytopes representing the identified archetypes for macrophages (A), fibroblasts (B), and endothelial cells (C) based on the Pareto optimality principle in PCA gene expression space. Each point corresponds to a cell, and distinct archetypes (AM1-AM5 for macrophages, AF1-AF5 for fibroblasts, and AE1-AE6 for endothelial cells) are highlighted. Ellipses at the archetypes represent their confidence intervals. The functions inferred for each archetype are listed. (D) Schematic illustrating the concept of universal versus tissue-specific archetypes. Archetypes with high diversity of tissues in their composition are considered universal, while those associated predominantly with a single tissue are tissue-specific. (E-G) Tissue composition of each archetype for macrophages (E), fibroblasts (F), and endothelial cells (G). Each bar represents an archetype, and different colors indicate tissue contributions. (H-J) Diversity analysis of archetypes using the Gini-Simpson index for macrophages (H), fibroblasts (I), and endothelial cells (J). A higher index indicates a more diverse, universal archetype, while a lower index suggests a tissue-specific archetype. (K-L) Visualization of fibroblast gene expression in t-SNE representation. Cells are colored by their tissue of origin (K) and by euclidean distance to their corresponding archetype (L, brighter shade represents closer distance to archetype and grey represents generalist cells).

### Supportive cell populations exhibit both universal and tissue-specific specializations

To explore whether supportive cell archetypes are shared across tissues or shaped by local context, we examined the tissue composition of cells located near each archetype in gene expression space. We defined these “proximal” cells using a proximity threshold, excluding generalist cells to focus on archetype-enriched regions (Methods). For each archetype, we quantified the diversity of its tissue representation using the Gini-Simpson index^51,52^, where values close to 1 indicate broad (universal) tissue distribution and values near 0 reflect tissue specificity (Fig. 2D, Methods).

Among macrophages, all five archetypes show high tissue diversity (Gini-Simpson > 0.8, Fig. 2E, H), suggesting that core macrophage functions, including immune signaling, phagocytosis, and homeostasis, are performed similarly across tissues. Some modest enrichments were present, such as fat-derived cells near the repair/homeostasis archetype (AM2) and skin-derived cells near the cytokine-production archetype (AM3).

In contrast, fibroblasts and endothelial cells display both universal and tissue-specific archetypes. Four fibroblast archetypes (AF1-AF3, AF5) are broadly shared across tissues, though each showed moderate enrichment (pancreas in AF1, large intestine in AF2, and thymus in AF3). The immune-signaling fibroblast archetype (AF4), however, was composed almost exclusively of skin fibroblasts (Gini-Simpson < 0.1, Fig. 2F, I). A similar pattern was observed in endothelial cells. While four archetypes (AE1, AE3, AE5-AE6) are broadly distributed across tissues, with enrichment from large intestine (AE3), lung (AE5), and skin (AE6), two archetypes (AE2 and AE4) are composed almost exclusively of cells from spleen and liver, respectively (Gini-Simpson = 0; Fig. 2G, J). These reflect known tissue-specific endothelial populations such as sinusoidal endothelium, which are uniquely adapted to the filtering and immune environments of the liver and spleen^53–56^. These universal and tissue-specific patterns are largely robust to variations in proximity thresholds (Fig. S2). One exception is the adaptive-immunity archetype of endothelial cells, AE5, which becomes lung-dominated at higher thresholds (Fig. S2O). To confirm that endothelial dominant tissue-specific archetypes from spleen and liver were not skewing our results, we repeated the analysis without these tissues. The core universal archetypes (AE1, AE3, AE6) remained intact, and additional specializations emerged for lung and skin endothelium related to immune regulation, barrier maintenance, coagulation regulation, and cell growth (Fig. S3).

Visualization of archetype assignments in t-SNE representation alongside tissue of origin further reveal how universal archetypes span multiple tissues, while tissue-specific archetypes capture localized specialization (Fig. 2K-L, Fig. S4, Methods). This illustrates how the Pareto framework complements clustering approaches: cells from different tissues performing similar tasks converge toward shared archetypes and are found in close or shared clusters, while cells from the same tissue can separate into distinct archetypes based on functional tradeoffs. Adjacent clusters that appear as continuous spectra reflect generalist cells positioned between archetypes (Fig. 2L).

Together, these results suggest that supportive cell types divide their labor both in conserved and context-specific manners. Skin fibroblasts emphasize immune signaling, consistent with constant exposure to external challenges, while liver and spleen endothelial cells specialize in permeability, endocytosis, innate immunity and mitochondrial energy production, aligning with their role in filtration and nutrient processing^57–59^. The coexistence of universal and locally tailored archetypes reflects a flexible division of labor that allows tissues to balance core supportive functions with specific physiological demands.

### A spectrum of archetype distributions reveals coordinated functional diversity across tissues

Following the identification of universal and tissue-specific archetypes, we next asked whether different tissues share similar division-of-labor patterns. To do so, we use an inverted perspective where we examine the distribution of archetypes in every tissue. For each cell type, we quantified the proportion of cells assigned to each archetype using the proximity analysis (Fig. 3A, Methods). This analysis revealed a continuum of division-of-labor strategies: some tissues contain a diverse mixture of archetypes, suggesting multitasking populations, while others are dominated by a single archetype, reflecting task specialization (Fig. 3B-D).

**Figure 3:**
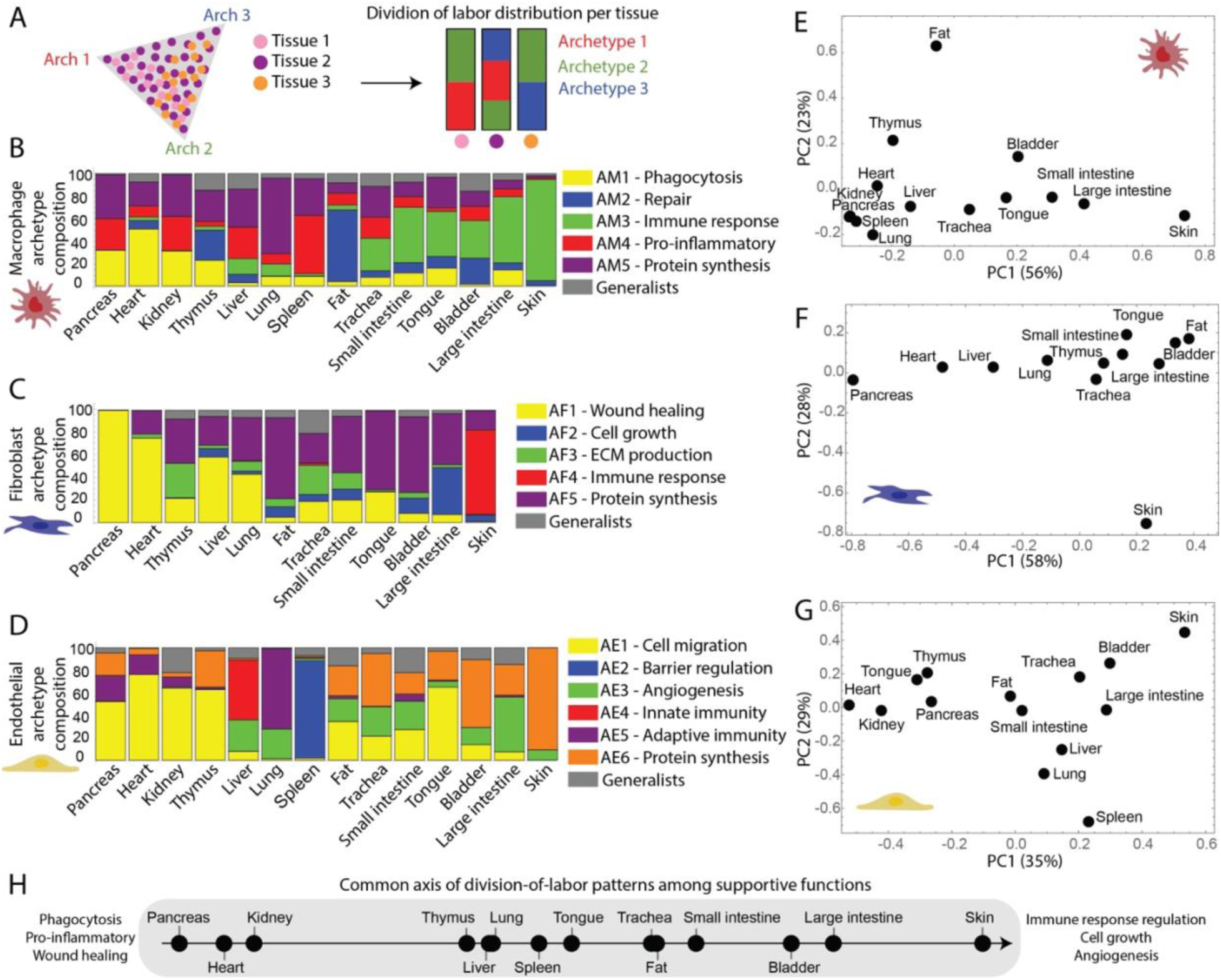
Supportive cells from different tissues show a variety of archetype profiles. (A) Schematic illustrating how archetype distributions are quantified per tissue. (B-D) Archetype composition analyses reveal the diversity of functional states among macrophages (B), fibroblasts (C), and endothelial cells (D) across various tissues. Each bar represents a specific tissue, with colors corresponding to different archetypes, indicating their relative contributions within each tissue. Generalists (in gray) are cells that did not cross the proximity threshold to any of the vertices. (E-G) Principal component analysis (PCA) of archetype composition for macrophages (E), fibroblasts (F), and endothelial cells (G) across different tissues. Each point corresponds to a tissue, positioned according to similarity in archetype composition. (H) Axis representing similarity in archetype distributions across tissues in all three cell types, from archetypes associated with phagocytosis, inflammation, and wound healing (left) to those involved in cell growth, angiogenesis, and immune regulation (right). The values were computed by taking the average of the first PC values in panels E-G.

Tissues such as the small intestine, liver, and trachea exhibit broad archetype diversity across all three supportive cell types, indicating a heterogeneous population structure with multiple functional roles coexisting. In contrast, other tissues display striking archetype polarization. For example, the skin consistently shows strong specialization across macrophages, fibroblasts, and endothelial cells, with each population dominated by a single archetype (Fig. 3B-D). This suggests that the skin microenvironment favors strongly differentiated functional roles, likely reflecting its constant exposure to external insults and need for barrier integrity. Other examples of skewed archetype composition include fat and lung macrophages, pancreatic and heart fibroblasts, and spleen endothelial cells. Each of these populations is dominated by a distinct archetype, suggesting highly tailored functional programs in these tissues.

We also examined the distribution of generalist cells, those located near the center of the archetype polytope performing a mixture of tasks (Fig. 3B-D). Generalists are more common in tissues with broad archetype diversity, such as the intestine or liver, and are underrepresented in tissues with polarized archetype profiles, such as skin and spleen. This implies that in multitasking tissues, cells may adopt hybrid strategies, whereas in specialized tissues, cells align more closely with distinct functional roles.

To better understand patterns of similarities between tissues in terms of supportive cell organization, we performed principal component analysis (PCA) on the archetype distributions across tissues (Fig. 3E-G, Fig. S5). This revealed a continuous distribution of tissues based on their supportive archetype configurations with a few divergent tissues. For example, skin and fat macrophages cluster separately from other tissues and from each other, reflecting distinct archetype biases (Fig. 3E). Skin fibroblasts also form an outlier (Fig. 3F), and spleen, lung, and liver endothelial cells separate from the rest (Fig. 3G), consistent with their unique functional demands.

Across all cell types, the first principal component captures a shared axis of variation that separates metabolically active tissues (e.g., liver, kidney, pancreas, heart) from mucosal or barrier tissues (e.g., large intestine, trachea, tongue, skin) (Fig. 3E-G). This suggests a fundamental tradeoff axis between metabolic support and structural/immune protection, reflected in the organization of supportive cell populations (summarized in Fig. 3H). This analysis reveals that supportive cell types adopt a diverse range of division-of-labor strategies across tissues, from flexible multitasking to rigid specialization, shaped by local physiological demands.

### Crosstalk among supportive cell archetypes reveals conserved and task-specific communication patterns

Fibroblasts, macrophages, and endothelial cells engage in extensive intercellular communication to coordinate tissue-level functions^60–62^. This communication is mediated by diffusible signals such as cytokines and growth factors, as well as indirect cues through ECM remodeling^9,63,64^. We next asked whether this signaling occurs broadly among all supportive cells or is restricted to specific functional subpopulations, and whether this signaling contributes to the organization of division of labor across and within cell types. To address this, we built an archetype-level communication network by integrating a curated database of ligand-receptor pairs^65^ with the transcriptional profiles of each archetype across the three cell populations. In this framework, a directed edge is drawn from archetype A to archetype B if archetype A expresses a ligand whose corresponding receptor is enriched in archetype B. For example, VEGFA expression in a macrophage archetype (AM3) and its receptor FLT1 in an endothelial archetype (AE3) indicates a potential directed signaling from AM3 to AE3 (Fig. 4A).

**Figure 4:**
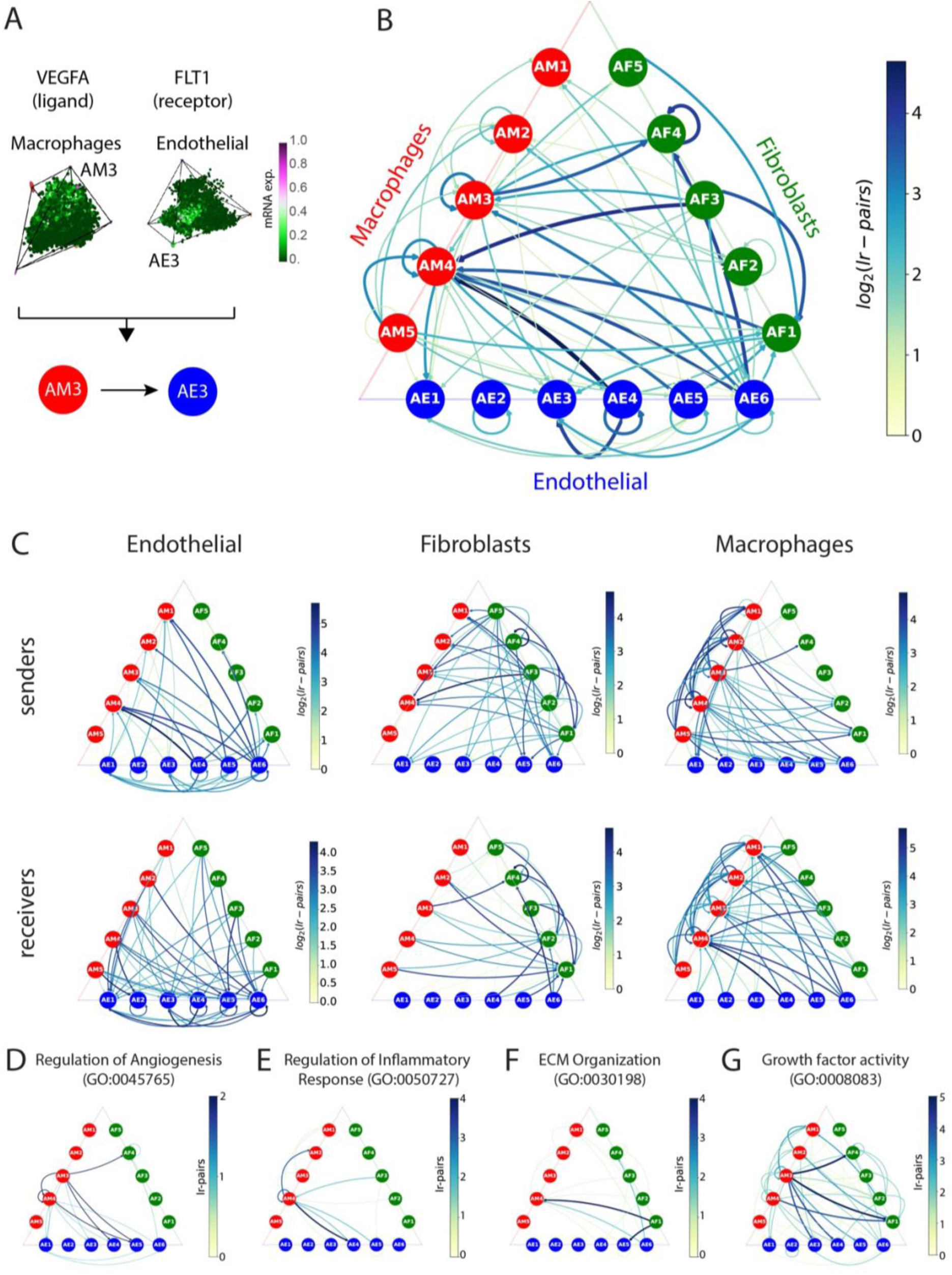
Archetype crosstalk networks within and among supportive cell populations. (A) Schematic illustrating the archetype-level ligand-receptor analysis. An archetype is considered to signal to another if a ligand enriched in the sender archetype has a corresponding receptor enriched in the receiver archetype. Example shown for VEGFA (ligand) enriched in macrophage archetype AM3 and its receptor FLT1 enriched in endothelial archetype AE3. (B) Global archetype crosstalk network integrating macrophage (red), fibroblast (green) and endothelial (blue) archetypes. Edge width and color reflect the number of ligand-receptor pairs (in log scale). Both within- and cross-cell-type links are shown; Analysis is restricted to archetypes with overlapping tissue composition to exclude edges from tissue segregation (Methods). (C) Ligand-receptor interactions for each cell type considered separately as senders (top) and receivers (bottom). Networks show the number of ligand-receptor pairs between archetypes. (D-G) Process-specific communication networks for archetypes focusing on regulation of angiogenesis (D), regulation of inflammatory response (E), ECM organization (F), and growth factor activity (G). For each process, only ligands and receptors that are included in the specified GO category were considered in the network.

We recently developed this approach to explore interactions among archetypes within the same cell type, and previously applied it to fibroblasts and macrophages from colon and lung tissues individually, revealing the potential regulatory role of cell communication in orchestrating division of labor^30^. Here, we extend this approach to capture interactions both within and across cell types, analyzing cells from multiple tissues simultaneously to identify communication patterns that are conserved across tissue contexts. To avoid spurious links driven by tissue segregation, we retain only interactions in which both ligand and receptor are enriched in archetypes with overlapping tissue composition (Methods). This analysis reveals dense interconnections both within and between cell types, indicating that cell-cell communication is closely coupled to functional specialization (Fig. 4B, Tables S4-S8).

Examining incoming and outgoing edges separately for each cell type reveals a pattern of hierarchy within the global network. Endothelial archetypes tend to signal outward, particularly to macrophages; fibroblast archetypes send signals primarily to macrophages and only sparsely to endothelial cells; and macrophages emerge as central hubs, receiving diverse inputs while also relaying signals broadly across the network (Fig. 4C). This communication topology is consistent with macrophages’ known roles as integrators of environmental information^5^.

To further explore whether these interactions reflect specific biological processes, we restricted our archetype-archetype signaling analysis to ligand-receptor pairs associated with angiogenesis, inflammatory regulation, and ECM organization. We found that these processes engage distinct subsets of archetypes, rather than being broadly distributed across entire cell populations. For example, angiogenesis-related signals are strongly enriched between subsets of endothelial and macrophage archetypes, with limited involvement of fibroblasts (Fig. 4D). In contrast, inflammation- and ECM-related communication include all three cell types, yet still show specificity, as only particular archetypes within each cell type participate in these interactions (Fig. 4E-F). These patterns of process-restricted communication that is restricted to subsets of archetypes supports the notion that intercellular communication is not merely a ubiquitous background process but rather an underlying driver of cellular specialization.

Notably, when restricting the analysis to ligand-receptor pairs associated with general growth factor activity, we observed a much broader pattern of communication (Fig. 4G). Nearly all archetypes across the three cell types participate in these interactions, suggesting that growth factor signaling serves as a more universal communication mode, potentially involved in basic cellular maintenance, survival, and proliferation^66,67^. In contrast, signaling associated with inflammatory responses, angiogenesis, or ECM remodeling is more targeted, and involves only specific subsets of archetypes that specialize in those functions. These process-specific interaction maps suggest that supportive cells divide labor not only through their intrinsic functions but also through their communication roles. Moreover, the observation that all three cell types participate in core, cross-tissue processes highlights, the tight regulatory coordination required to maintain tissue homeostasis across diverse environments.

### Universal archetypes are recapitulated in analysis of individual tissues

To assess the robustness and context-dependence of the pan-tissue archetypes identified across combined datasets (Fig. 2), we repeated the Pareto analysis separately for each tissue out of the 14, and for each cell type (macrophages, fibroblasts, and endothelial cells). This within-tissue analysis allows us to evaluate whether the universal archetypes also emerge when tissues are analyzed individually, and whether additional tissue-specific specializations can be detected. In Figure 5, we present the results for fibroblasts, while parallel analyses for macrophages and endothelial cells are shown in Figures S7-S8.

**Figure 5:**
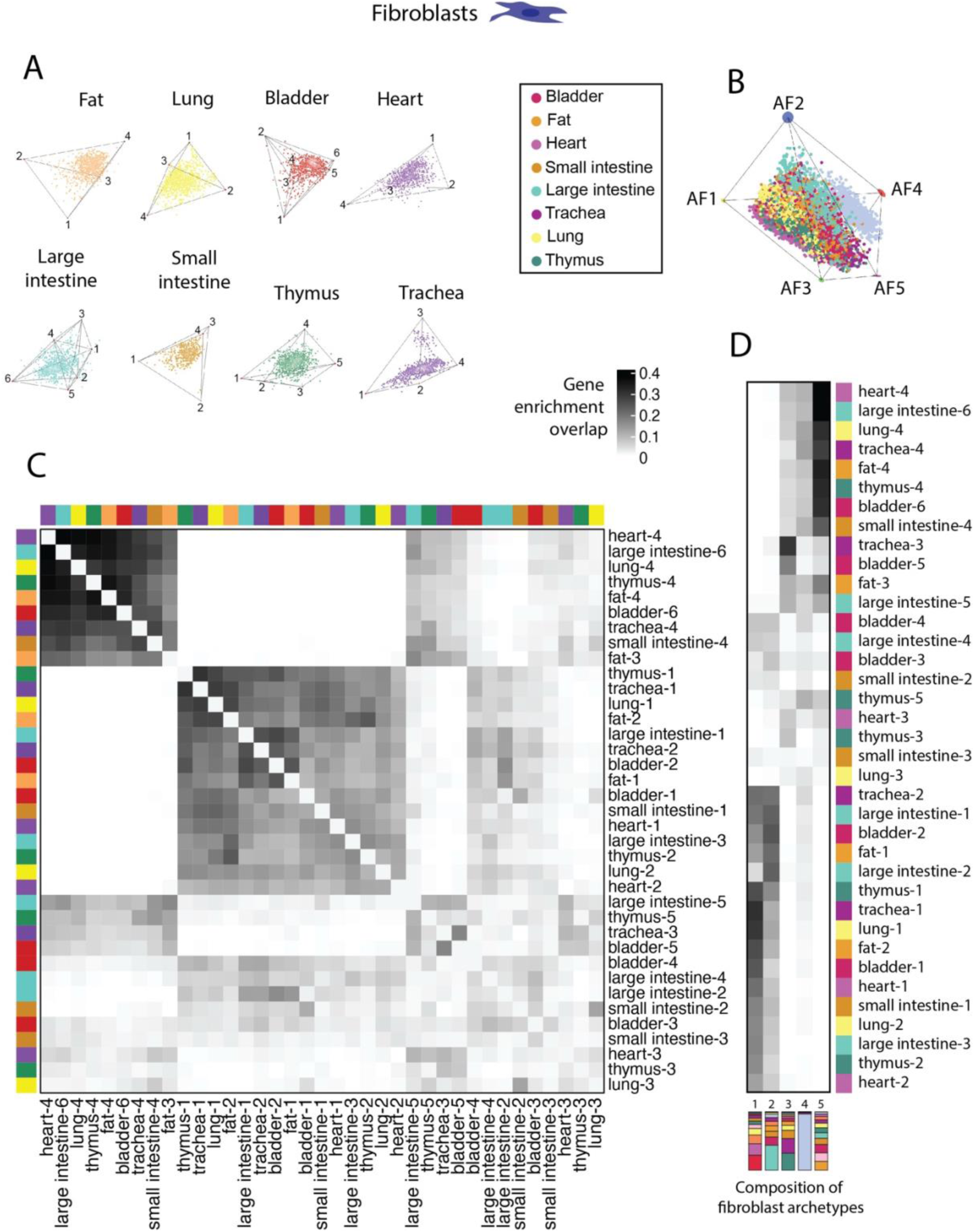
Universal archetype robustness and tissue-specific organization of fibroblasts. (A) Archetype polytopes for fibroblasts from individual tissues, with each cell colored by tissue of origin. Numbers indicate the archetypes identified within each tissue. Only tissues with significant p-values are presented (Table S9). (B) Global polytope of pan-tissue fibroblast archetypes derived when all tissues combined (as in Fig. 2B), shown here to illustrate how individual fibroblast populations separate by tissue within the shared archetype space. (C) Heatmap comparing enriched-gene overlaps among all within-tissue fibroblast archetypes across tissues. Row clustering is based on the similarity of enriched gene sets between archetypes. Darker shading indicates higher Jaccard overlap. Tissue of origin is indicated by the color bars along both axes. This comparison highlights how fibroblast archetypes from different tissues share or diverge in their gene enrichment profiles. (D) Heatmap showing the overlap in enriched genes between individual-tissue fibroblast archetypes (rows) and pan-tissue fibroblast archetypes (columns). Rows are clustered according to their similarity to the global archetypes. Darker shading indicates higher Jaccard overlap, with tissue origin for each archetype annotated by the color bar.

We find that in the majority of the tissues, gene expression profiles are well-described by polytopes with 4-6 archetypes (64% in macrophages and endothelial cells, and 67% in fibroblasts; Fig. 5A, Fig. S6, Table S9, Methods), consistent with the presence of functional tradeoffs and division of labor at the tissue level. One notable exception is the skin, where fibroblasts, macrophages, and endothelial cells show poor polytope fits, aligning with our previous observations of dominant, task-specific specializations and minimal tradeoff-driven heterogeneity.

To compare the within-tissue archetypes (Fig. 5A) with the pan-tissue ones (Fig. 5B), we calculated gene set similarities using the Jaccard index between the enriched gene sets of each pair of archetypes (Methods). This comparison reveals that every tissue includes multiple within-tissue archetypes that show high similarity to universal pan-tissue archetypes (Fig. 5D), demonstrating that conserved functional programs are preserved even when tissues are analyzed in isolation. Furthermore, within-tissue archetypes that map to the same universal pan-tissue archetype are themselves highly similar across tissues (Fig. 5C), reinforcing the notion that they represent shared, conserved functional roles.

At the same time, several within-tissue archetypes do not correspond to any of the pan-tissue archetypes, highlighting additional tissue-specific specializations that become apparent only in the context of individual tissue analysis. For example, bladder, spleen, and lung macrophages exhibit distinct mitochondrial and metabolic programs (Fig. S7), and endothelial cells from the large intestine display enrichment in contractile functions (Fig. S8). Importantly, pan-tissue archetypes that were identified as tissue-specific, such as the spleen- and liver-enriched endothelial archetypes (AE2 and AE4), closely match the archetypes independently inferred in those same tissues, validating the identity of these specialized programs (Fig. S8A). Together, this analysis supports a principle in which supportive cell populations in each tissue adopt multiple functional specializations. Some of these archetypes are universal and conserved across tissues, while others are uniquely tailored to local tissue context.

### Conservation and divergence of supportive cell archetypes across species

To explore the evolutionary conservation of division of labor among supportive cell types, we applied our archetype inference framework to scRNA-seq data of macrophages, fibroblasts, and endothelial cells from 13 murine tissues (data from^46,68,69^). Using the same Pareto Task Inference approach applied in human samples, we identified five significant archetypes per cell type (p < 0.001, Fig. 6A-C, Methods), consistent with the presence of tradeoffs among distinct functional programs in the mouse as well.

**Figure 6:**
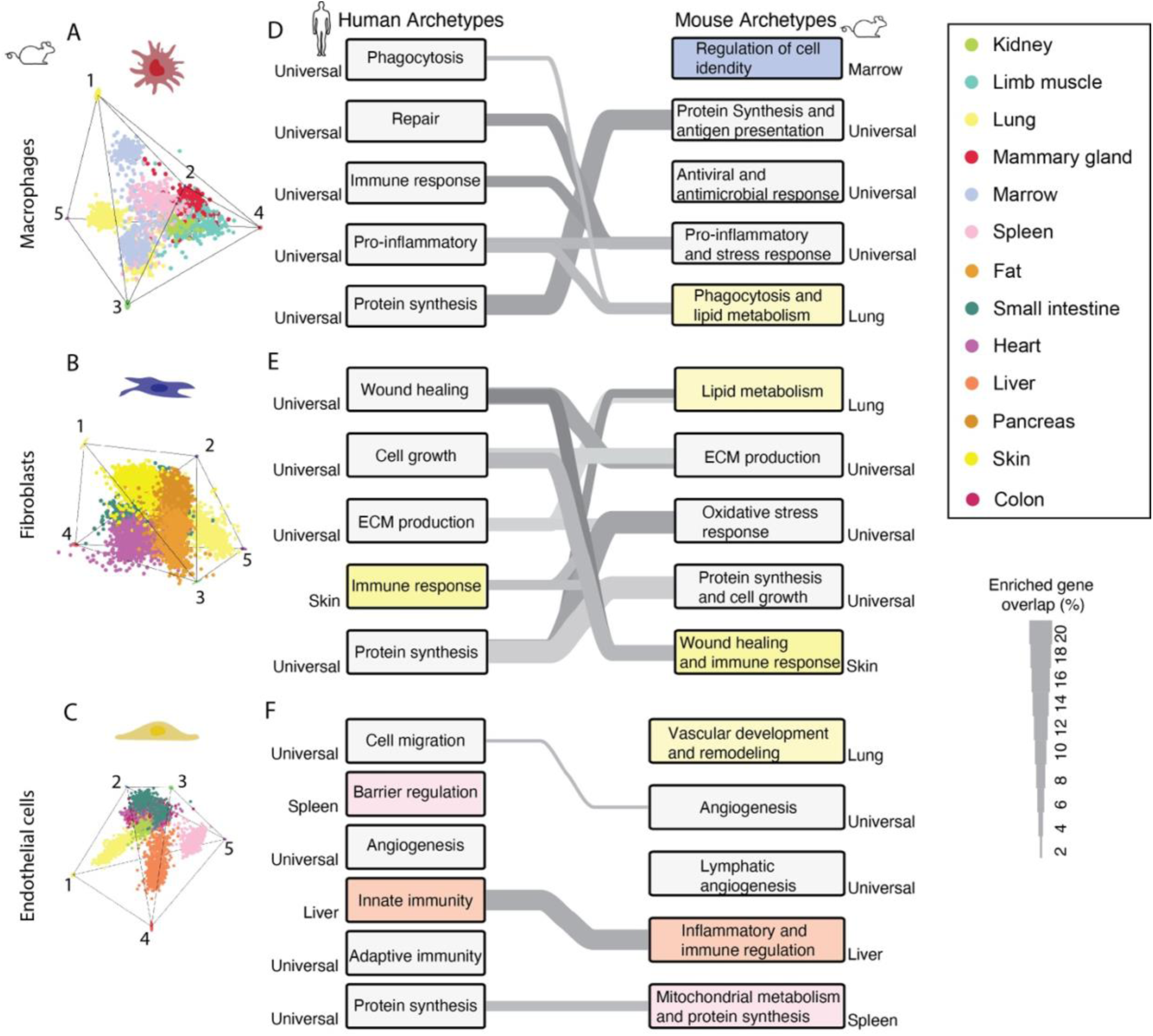
Comparison between human and murine archetypes. (A-C) Polytopes illustrating the identified archetypes for macrophages (A), fibroblasts (B), and endothelial cells (C) sampled from 13 different murine tissues. Each point corresponds to a cell, with different colors representing different tissue of origin. (D-F) The inferred human (left) and murine (right) archetype functionality for macrophage (D), fibroblast (E), and endothelial cells (F). Tissue-specific archetypes are colored according to the tissue color code and universal archetypes are colored in light gray. Gray lines connecting between human and murine archetypes represent statistically significant overlaps between enriched gene sets of each archetype, with line thickness corresponding to the percentage of overlap.

To quantify correspondence between species, we compared gene enrichment patterns between mouse and human archetypes, identifying functionally matched archetypes across species (Tables S10-S12). To match archetypes, we only retained overlaps that were statistically significant based on a shuffled distribution of gene enrichment across archetypes (Fig. S9A-C, Methods). We find that several universal archetypes identified in humans have clear counterparts in the mouse dataset. For example, macrophage archetypes associated with phagocytosis, pro-inflammatory response, and protein synthesis showed strong cross-species correspondence (Fig. 6D), as did fibroblast archetypes specializing in ECM production, wound healing, and cell growth (Fig. 6E). Endothelial archetypes show more limited overlap with archetypes linked to innate immunity, angiogenesis, and biosynthesis processes exhibiting conserved transcriptional signatures between humans and mice (Fig. 6F).

Tissue-specific archetypes were also largely conserved. For example, endothelial archetypes restricted to the liver and spleen appear also in the murine data as well as skin-specific archetype in mouse fibroblasts. This indicates that both the universal and tissue-specific architecture of supportive cell function is broadly maintained between humans and mice. Two notable exceptions highlight species- or dataset-specific divergence. First, we identified a lung-specific macrophage archetype in the mouse that overlaps with a universal human macrophage archetype. This human archetype is enriched in lung macrophages, suggesting that its tissue association is more narrowly defined in mice. Second, a bone marrow-specific macrophage archetype emerged in the mouse dataset, which has no counterpart in the human data, likely reflecting the fact that bone marrow was not included in the human tissue panel (Fig. 6D). This murine archetype captures functions unique to marrow-resident macrophages, which differ in role and origin from other tissue-resident macrophage populations^70,71^.

Despite the strong functional correspondences, the overall gene-level overlap between matched archetypes remained modest relative to the substantial overlap in highly variable genes between species. This suggests that the divergence in archetype structure is not simply due to technical or gene set differences. To further probe this, we co-embedded matched human and mouse tissues in a shared low-dimensional expression space (Fig. S9D-F). The first principal component consistently separated cells by species, indicating a dominant organismal effect, whereas the second axis captured tissue-specific differences. Additionally, mouse cells are clustered closely together, while human cells are more broadly dispersed. This pattern may emerge from the biological variability inherent in human samples, which span a diverse range of donors, ages, genetic backgrounds, metabolic states, and immune histories. In contrast, the mouse data derive from genetically identical, young adult animals housed in controlled laboratory conditions. These differences likely contribute to the broader, more continuous archetype distributions observed in humans and to the emergence of additional archetypes reflecting population-level diversity.

Overall, these findings suggest that while the distribution of tradeoffs among supportive cell functions is largely conserved across species, the specific transcriptional implementation of these programs is shaped by organism-specific and contextual factors. This highlights the importance of incorporating human-specific data to fully capture the spectrum of supportive cell states relevant to tissue homeostasis and disease.

## Discussion

In this study, we systematically mapped patterns of division of labor among three supportive cell populations that are found in most mammalian tissues: macrophages, fibroblasts, and capillary endothelial cells, across 14 human tissues, and compared them to murine counterparts. Using the Pareto optimality framework, we identified archetypes that represent transcriptional programs specialized for distinct functions. These archetypes revealed both universal patterns shared across tissues and specialized states shaped by local microenvironments. Our analysis further uncovers universal tradeoffs between core supportive functions such as immune regulation, wound healing, angiogenesis, ECM remodeling, and metabolic support, that are required for tissue homeostasis and recur across diverse tissues. Communication analysis further revealed that distinct archetypes engage in specialized ligand-receptor signaling, suggesting that division of labor is not only intrinsic but also actively coordinated between cell types. These findings offer a unified cross-tissue perspective on supportive cell organization and the principles underlying functional specialization and intercellular coordination, highlighting both conserved organizational principles and tissue-specific deviations and providing a systems-level view of supportive cell ecology in human tissues.

Our results extend and complement previous single-cell studies that identified shared clusters within each supportive cell type. In endothelial cells, for example, cross-tissue analyses revealed both angiotypic signatures and tissue-specific vascular profiles across 19 organs^39^. Similar efforts in myeloid cells identified conserved *S100A8*/*S100A12*-high classical monocytes and *TREM2*⁺ macrophages, alongside tissue-specific states such as *PPARG*⁺/*MARCO*⁺ alveolar macrophages in the lung and *CLEC4F*⁺/*VSIG4*⁺ Kupffer cells in the liver^40^. Related work has shown that shared immune programs are further shaped by inhibitory receptor networks that impose context-dependent thresholds and feedback control^72,73^. In fibroblasts, mouse studies have identified both *Col15a1*⁺ ECM-organizing fibroblasts shared across tissues and more specialized subtypes^46^. Our work agrees with these findings and provides a new perspective by framing them within a tradeoff structure. Many of these previously described clusters correspond to universal archetypes in our analysis: *S100A8/S100A12* are enriched in AM4 (a pro-inflammatory universal macrophage archetype), *MARCO* in AM2 (prominent in adipose tissue but shared across tissues), and *COL15A1* in AF1 (a universal ECM-production fibroblast archetype). This tradeoff framework reveals how universal states arise from balancing competing demands and how tissue-specific pressures lead to tailored adaptations.

This perspective also clarifies how supportive cell division of labor is organized across tissues. We identified a common cross-tissue axis that spans metabolically active tissues (e.g., liver, pancreas, heart) to barrier tissues (e.g., skin, colon, trachea), along which macrophages, fibroblasts, and endothelial cells align in their archetypal distributions. Tissues at opposite ends of this axis show striking contrasts in specialization. For example, in the skin, each supportive cell type primarily occupies a single archetype, suggesting highly constrained functional roles, whereas tissues like liver and lung exhibit broader archetype diversity, reflecting a wider functional repertoire. This pattern highlights the adaptability of supportive cells and their capacity to tune function to the needs of each tissue. In barrier tissues, macrophages favor immune-regulatory states that minimize collateral damage, while internal tissues favor pro-inflammatory and reparative states. Similarly, fibroblasts and endothelial cells in energy-demanding tissues tend to adopt metabolically active archetypes.

These findings have implications for understanding tissue vulnerability and disease. Pancreatic fibroblasts, for example, are concentrated near a single wound-healing archetype distinct from other tissues, possibly explaining their pronounced role in pancreatic ductal adenocarcinoma, where they support ECM remodeling, immune evasion, and metabolic crosstalk with tumor cells^74–76^. More broadly, deviations from typical division-of-labor patterns, such as overrepresentation of matrix-producing fibroblasts in fibrosis or inflammation-associated subtypes in chronic disease^77,78^, may reflect a breakdown in the normal archetypal balance. Tissues that display a skewed archetype distribution, deviating from the common division-of-labor patterns observed across most tissues, may be less resilient to perturbations. Such reduced flexibility could increase their vulnerability to environmental or pathological challenges and predispose them to disease.

Our intercellular communication analysis further supports the existence of coordinated division of labor. Universal archetypes engage in conserved ligand-receptor interactions across tissues, making them strong candidates for core regulatory pathways that coordinate supportive cell function. Process-specific communication networks showing that distinct archetypes mediate angiogenesis, immune regulation, and ECM remodeling further support the idea that division of labor is regulated through specialized intercellular signaling. Comparing these communication networks in healthy versus diseased tissues may identify key pathways whose disruption drives pathological remodeling.

Cross-species comparisons show that the overall distribution of universal and tissue-specific archetypes is conserved, including skin fibroblast specialization and liver/spleen endothelial programs. However, mouse archetypes tend to be more tissue-specific. For example, a lung-specific macrophage archetype in mice corresponds to a universal human one. These differences may be explained by the reduced biological variability in laboratory mice compared to the genetically and physiologically diverse human cohorts^79^. Nonetheless, the recurrence of core archetypes in both species suggests that the fundamental tradeoffs governing supportive cell specialization are evolutionarily conserved and necessary for tissue homeostasis.

Looking ahead, a key question is how supportive archetypes integrate with the functions of parenchymal cells in each tissue, and how these relationships are altered in disease. In cancer or fibrosis, novel archetypes may emerge that support pathological tissue states, such as persistent ECM production or immunosuppression. A particularly informative direction will be perturbation experiments targeting transcription factors enriched in archetypes that resemble putative disease precursors, to test whether modulating these regulatory nodes can reprogram cells and restore healthy archetype balance. Such longitudinal or perturbation-based studies will be critical to determine whether shifts in archetype composition are causal or consequential in disease progression, and whether restoring archetypal balance can help reestablish healthy tissue organization and function.

## Acknowledgements

We thank Ruslan Medzhitov, Xu Zhou, Uri Alon, Avi Mayo, Shoval Miyara and members of the Adler lab for meaningful discussions. This study was supported by the Israel Science Foundation (ISF, Grant No. 3372/24) and the United States-Israel Binational Science Foundation (BSF, Grant No. 2023231). MA is the recipient of the Alon Fellowship, Israel Council for Higher Education.

## Methods

### Analysis of single-cell RNA sequencing across 14 human tissues

We considered cells from the following tissues: heart (including cardiac related tissue), lung, fat (including adipose tissue, subcutaneous adipose tissue), thymus, pancreas, skin, liver, spleen, tongue, trachea, large intestine (or colon), small intestine (including duodenum, jejunum, ileum), bladder and kidney. The choice of tissues was based on availability of data and minimal number of cells for further analysis. In total, we analyzed 11,749 macrophages, 9,899 fibroblasts, and 10,207 endothelial cells.

For fibroblasts - we analyzed data from^47^. If more than 1,000 cells were available from a specific tissue, 1,000 cells were chosen randomly, if a tissue had fewer than 1,000 cells, we included all available cells. This led to raw counts from a total of 9,899 cells: 1,000 cells from the large intestine, bladder, fat, tongue, heart, skin, trachea, lung, thymus; 544 cells from small intestine, 194 from pancreas and 161 from liver. Each cell’s expression profile was normalized to its total counts, followed by a log1p transformation. We then filtered genes based on their mean and standard deviation, using thresholds of: mean threshold = 1e-5 and std threshold = 5e-6. After filtering the genes, z-score normalization was applied, resulting in a final set of 10,118 genes.

For endothelial cells - we analyzed data from^47^ included endothelial cells in the analysis if they originated from one of the selected tissues above and were classified as one of the following cell types: capillary endothelial cell, cardiac endothelial cell, colon endothelial cell, endothelial cell, endothelial cell of arteriole or endothelial cell of venule. If more than 1000 cells were available from a specific tissue, 1000 cells were chosen randomly, leading to raw counts from a total of 10207 cells: 1000 cells from heart, lung, fat, thymus, pancreas, skin, liver, spleen, and tongue; 549 from the trachea; 256 from the large intestine; 177 from the small intestine; 124 from the bladder and 101 cells from the kidney. We normalized the data as described for fibroblasts. Highly variable genes were selected by taking the 8,685 genes with mean expression>1.5e-5 and standard deviation>5e-5. Finally, z-score normalization was applied.

For macrophages - we download data from^47^. Cells were included if they originated from one of the selected tissues and were classified as monocyte or macrophage populations. If more than 1000 cells were available from a specific tissue, 1000 cells were chosen randomly, leading to raw counts from a total of 11,748 cells: 1,000 cells from the bladder, fat, liver, skin, pancreas, lung, and spleen; 815 from heart; 814 from trachea; 796 from thymus; 748 from large intestine; 709 from tongue; 444 from small intestine; and 422 from kidney. We normalized the data as described for fibroblasts and endothelial cells. We considered the 8,159 genes with mean expression > 9.5 × 10⁻⁶ and standard deviation > 5 × 10⁻⁵, after which z-score normalization was applied. All preprocessing and normalization steps were performed in Python using Scanpy and AnnData.

### Applying the Pareto Archetype Inference to the single-cell data

We inferred division of labor patterns between biological function for each supportive cell population individually by applying the ParTI MATLAB package^50^ on the normalized gene expression data using the following parameters: the PCHA algorithm was applied to find the simplex, Explained Variance was calculated up to the 8th dimension, and the bin size for feature enrichments was set to 0.05. The number of archetypes for each cell population was chosen according to the value suggested by the algorithm, based on the elbow in the explained variance plot. Once the algorithm finds a simplex that fits the data, the software calculates the ratio between the simplex volume and the convex hull. A p-value for the fitted simplex is provided by randomly shuffling the data points in PCA space and recalculating the volumes ratio for 1000 times.

### Gene enrichment analysis

To interpret the biological functions associated with each archetype identified by ParTI, we performed a gene enrichment analysis on genes enriched near the archetypes in gene expression space. ParTI identifies archetypes and computes for each gene its mean gene expression across 20 bins organized by their proximity to these archetypes. Genes that are strongly associated with a specific archetype (i.e., expressed most highly in cells near that archetype) are considered enriched near that archetype. For each archetype, we selected the top ranking enriched genes based on their adjusted p-values and effect size in gene enrichment, and characterized their biological functions using a combination of automated Gene Ontology (GO) term enrichment analysis and manual annotation based on GeneCards^80^ database descriptions. This combined approach enabled us to assign functional labels to each archetype, based on robust patterns of gene enrichment and known biological roles of the associated genes.

Examples of characteristic enriched genes:

Macrophage archetypes:
(AM1) MERTK, ELMO1, DOCK8, JAK2, NFATC3, and MAP2K5
(AM2) LYVE1, CD163, and ADAM17
(AM3) CXCL8 and HLA-DRA
(AM4) NFKB1, TREM1, and LYZ
(AM5) Protein synthesis genes

Fibroblast archetypes:
(AF1) COL15A1, COL4A1, COL4A4, and LAMA2
(AF2) PFKFB3, GLS, NFAT5, STAT3, ETS2, SOX5, FOXO3, ASXL1, CHD1, and SIRT1
(AF3) DCN, FBLN1, LUM, ACTB, MYL12A, and MYL6
(AF4) IL15RA, IL32, IL1R1, IL6, CXCL1/2/3/8/14
(AF5) Protein synthesis and mitochondrial genes

Endothelial archetypes:
(AE1) PTK2 and DOCK1
(AE2) CDH11, CTNND2, and STAB2
(AE3) GLS, NAMPT, and PPARD
(AE4) MT-ND4L, FGA/B/G, APOA1, F2R, and CXCL16
(AE5) MHC class I (HLA-A/B/C/E/F) and class II (HLA-DRB1, HLA-DPA1) genes
(AE6) CDK2AP1 and HMGA1

### Analysis of tissue composition of archetypes

To characterize the tissue composition of every archetype, we mapped each cell to the archetype it is closest to in the original gene expression space. Each cell was represented as a vector inside the polytope defined by the archetype vertices, and its position was expressed as a weighted sum of the archetype vectors. The archetype with the largest weight was taken as the archetype the cell is assigned to. To filter out generalist cells and focus on those with strong archetype affinity, we applied a proximity threshold: cells whose maximum weight was ≥ 0.30 were assigned to their archetype, whereas cells below that threshold were labelled as generalists. We note that for different numbers of archetypes, this threshold may vary. For each archetype, we counted the assigned cells from every tissue and calculated the proportional representation of each tissue within that archetype. We used this archetype assignment of each cell to generate Figure S4 as well.

### Quantifying archetype diversity

To quantify diversity of archetype composition, we calculated the normalized Gini-Simpson index (𝐺𝑆) for each archetype based on the following equation: 𝐺𝑆 = (1 − ∑_𝑖_(𝑝_𝑖_^2^))/𝐺𝑆_𝑚𝑎𝑥_. Where 𝑝_𝑖_ - is the proportion of tissue *i* near the archetype we consider, and 𝐺𝑆_𝑚𝑎𝑥_ is the maximal value the diversity index can be given the number of tissues considered. For the human data, we chose a threshold of 𝐺𝑆 = 0.2 to define specificity of archetypes, where archetypes with 𝐺𝑆 < 0.2 are considered tissue specific and archetypes with 𝐺𝑆 > 0.2 are defined as universal (common to diverse tissues). In the mouse data, since we analyzed fewer tissues per cell type, we considered a higher threshold for diversity of 𝐺𝑆 = 0.4.

### Analysis of cross-tissue endothelial archetypes without the spleen and the liver

Starting from the raw endothelial cells counts, we removed all cells originated from the liver or spleen, and normalized and filtered the data as described for the full tissue set (see Analysis of single-cell sequencing across 14 human tissues) leading to a gene expression matrix of 8514 genes and 8207 cells. The ParTI package suggested a simplex with 5 archetypes (p-value<0.001).

### Analysis of archetype composition across tissues

To compare archetype composition of every cell type population across tissues, we applied principal component analysis (PCA) on the archetype distribution where data points represent the different tissues, and the axes are the archetype composition (based on the assignment of each cell to the archetype it is closest to, without generalist cells).

### Analysis of ligand-receptor interactions among archetypes

To construct the archetype-level cell-cell communication network, we combined a curated ligand-receptor database^65^ with gene enrichment profiles identified across all archetypes. We retained ligand-receptor pairs only when the ligand was enriched in one archetype and its corresponding receptor was enriched in another. To ensure these interactions were not driven solely by tissue-specific expression, we further filtered the interactions based on the similarity in tissue composition between the archetypes: only pairs whose tissue enrichment profiles exceeded a cosine similarity threshold of 0.1 were considered. In addition, we required that each enriched gene pass a minimum threshold for expression specificity, quantified by the ParTI algorithm’s “mean difference” metric, which captures how distinct the gene’s expression is in cells closest to a given archetype versus those furthest away. Only genes with a mean difference above 0.3 were included. For pathway-specific interaction maps (e.g., angiogenesis, inflammation, ECM organization), the ligand-receptor dataset was further restricted to genes associated with the relevant Gene Ontology (GO) term, and only interactions where both ligand and receptor matched the GO category were retained.

### Pareto archetype inference to supportive cells from each tissue individually

To investigate within-tissue functional programs in human supportive cells, we applied ParTi^50^ separately to macrophages, fibroblasts, and endothelial cells from each tissue. For each tissue - cell type combination, we first subset the relevant cells and performed several preprocessing steps. Gene expression matrices were normalized by row sums to account for total expression per cell, and lowly expressed genes (mean expression below 5 ∗ 10^−7^ were filtered out. The data were then log-transformed to stabilize variance, and highly variable genes (HVGs) were selected based on their average expression and standard deviation. Specifically, we retained genes whose mean and standard deviation exceeded the 55th percentile of the respective distributions. To control for donor-level variation, we applied a z-score transformation separately for each donor, standardizing gene expression values within each donor group by subtracting the mean and dividing by the standard deviation. Following preprocessing, we applied the ParTI algorithm to each normalized dataset to identify the low-dimensional Pareto front (polytope). The number of archetypes was initially selected based on ParTI’s recommendations, but final decisions were made based on the combination of statistical significance (p-values) and the stability of the archetype structure (standard deviation of archetype coordinates). If the recommended solution had a p-value greater than 0.05 or high variability, we tested alternative models with 4-6 archetypes and chose the best-supported configuration. These comparisons and selection criteria are summarized in Table S9. For each inferred polytope, we identified genes enriched near each archetype. To quantify the similarity between within-tissue and pan-tissue archetypes (defined from pooled data across all tissues), we computed the Jaccard index between enriched gene sets. For each archetype from an individual tissue analysis, we calculated the Jaccard similarity with each of the pan-tissue archetypes from the same cell type, defined as the size of the intersection divided by the size of the union of the two gene sets.

### t-SNE visualization of distance to archetype

t-Distributed Stochastic Neighbor Embedding (t-SNE) was calculated on the normalized gene expression matrix of each cell type with the MATLAB t-SNE function (Algorithm = exact, Distance = cosine). Cells were assigned to their closest archetype if their corresponding maximum weight was at least 0.3.

### Analysis of single-cell RNA sequencing across 13 mouse tissues

For fibroblasts - we downloaded data from^46^ If more than 1000 cells were available from a specific tissue, 1000 cells were chosen randomly, leading to raw counts from a total of 6000 cells: 1,000 cells from the intestine, fat, heart, skin, pancreas, lung. For each cell, we normalized the expression values to its total counts. We then applied a log1p transformation, then we filtered out genes with mean expression smaller than 8e-7 and standard deviation below 5e-5. Finally we performed z-score normalization across tissues. After these preprocessing steps, 8,452 genes remained.

For endothelial cells - we downloaded murine endothelial single cell RNA-seq data from Kalucka et al. We considered the following tissues: colon (741 cells), heart (485), kidney (245), liver (705), lung (588), small intestine (452), and spleen (750). 7173 genes with a mean expression> 8e-6 and standard deviation>7e-5 were selected.

For macrophages - we downloaded droplet (10×) macrophage data from Tabula Muris^68^. We kept only cells labeled as macrophage, monocyte, alveolar macrophage, classical monocyte, or non-classical monocyte. After filtering, 2,568 cells remained: 748 from marrow, 724 from lung, 464 from spleen, 307 from limb muscle, 186 from mammary gland, and 139 from kidney. We normalized the counts as described earlier, selected 8,210 highly variable genes (mean > 6.5e - 6; SD > 3.7e-5), and then applied z-score scaling.

### Statistical significance analysis of the comparison of gene enrichment between human and mouse archetypes

To assess the significance of the overlap between enriched genes in human and mouse archetypes, we performed a permutation test. Specifically, for each mouse archetype, we took the list of enriched genes and randomly shuffled the gene names. We then calculated the percentage of overlap between these shuffled gene sets and the corresponding enriched gene sets in humans. This process was repeated 10,000 times to generate a null distribution. The p-value was computed as the proportion of permutations where the overlap exceeded that of the original (non-shuffled) gene sets. To adjust for multiple comparisons within each cell type, we applied the Benjamini-Hochberg false discovery rate (FDR) procedure. Q-values were computed using the ‘mafdr’ function in MATLAB. A q-value below 0.05 was considered statistically significant.

### Comparison of mouse and human tissue gene expression profiles

To determine whether species or tissue type has a dominant influence on the differences in archetypes between mouse and human, we compared their gene expression profiles. For this analysis, we selected two tissues shared by both species for each cell type. We standardized the gene set by identifying orthologs common to mice and humans, and applied the same normalization process used in the main analysis. We performed principal component analysis (PCA) on the data from both species and separately for each cell type, using expression levels of the most highly variable shared genes.

## Supplementary Figures

**Figure S1:**
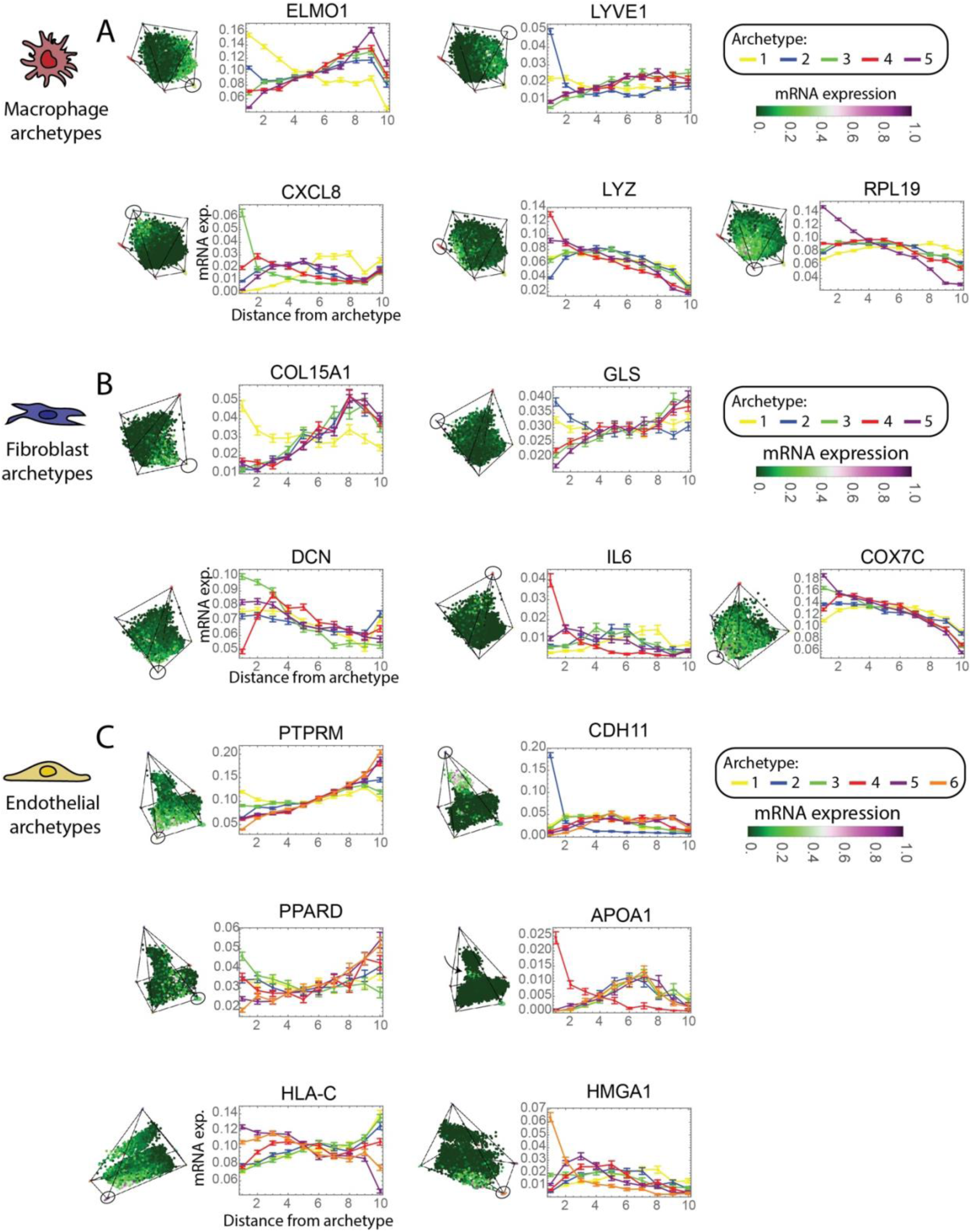
Enrichment profiles of selected genes across archetypes in each supportive cell population. Gene expression gradients are shown for selected genes across archetypes of (A) macrophages, (B) fibroblasts, and (C) endothelial cells. Each panel presents a 3D PCA of gene expression space (left) and a corresponding line plot (right) depicting mRNA expression levels as a function of distance from the indicated archetype. Curves represent the average gene expression in every bin of increasing distance from each archetype and STD are marked in each bin. Each color corresponds to a different archetype.

**Figure S2:**
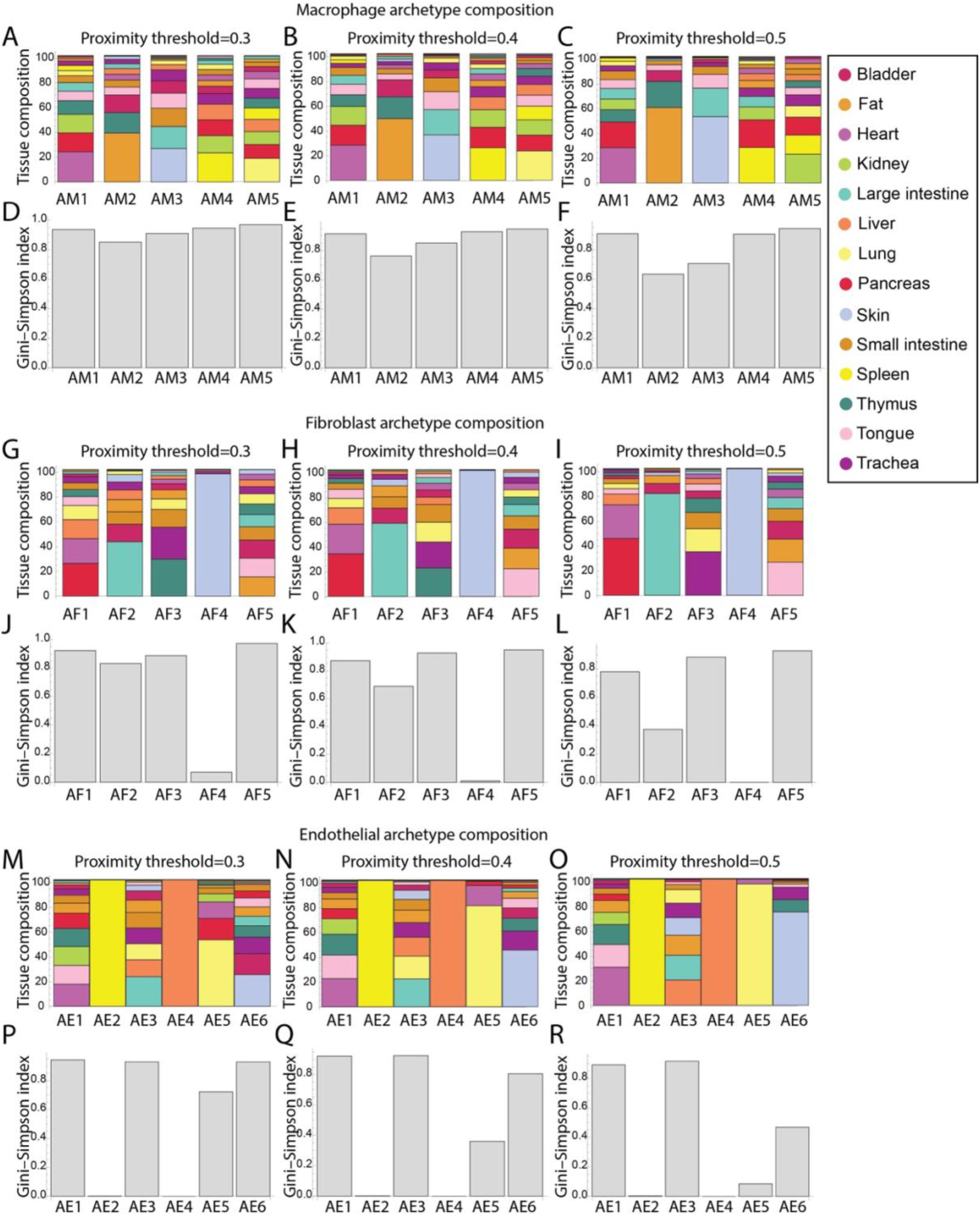
Effect of proximity threshold on tissue composition and diversity of supportive-cell archetypes. (A-C) Tissue composition of macrophage archetypes at thresholds 0.3, 0.4, 0.5, higher thresholds increase tissue specificity. (D-F) Corresponding Gini-Simpson indices, diversity decreases with threshold. A similar pattern is observed for fibroblast (G-I, J-L) and endothelial (M-O, P-R) archetypes.

**Figure S3:**
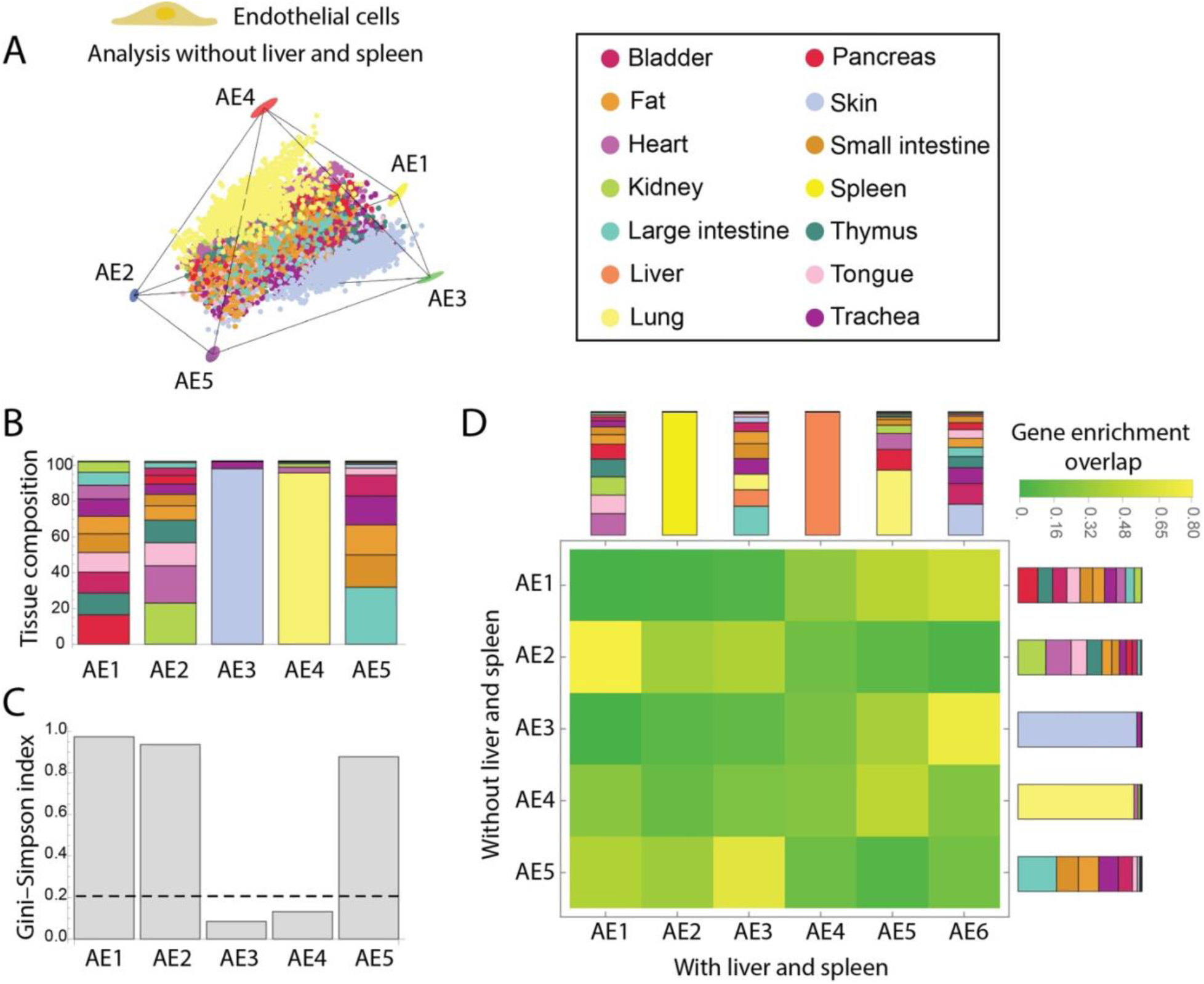
Endothelial cell archetypes analysis without liver and spleen. (A) Polytope representing the identified archetypes for endothelial cells from all tissues except for the liver and the spleen. Each point corresponds to an endothelial cell, and the five identified archetypes (AE1-AE5) are highlighted. Ellipses at the archetypes represent their confidence intervals. (B) Tissue composition of each archetype. Each bar represents an archetype, and different colors indicate tissue relative contributions. (C) Archetypes diversity analysis using the Gini-Simpson Index. The dashed line represents a threshold between universal archetypes (high index) and tissue-specific archetypes (low index). (D) The overlap between enriched genes in each archetype inferred when considering all tissues (columns, Fig. 2C) and archetypes inferred when excluding the liver and the spleen (rows). Bars represent archetype tissue composition.

**Figure S4:**
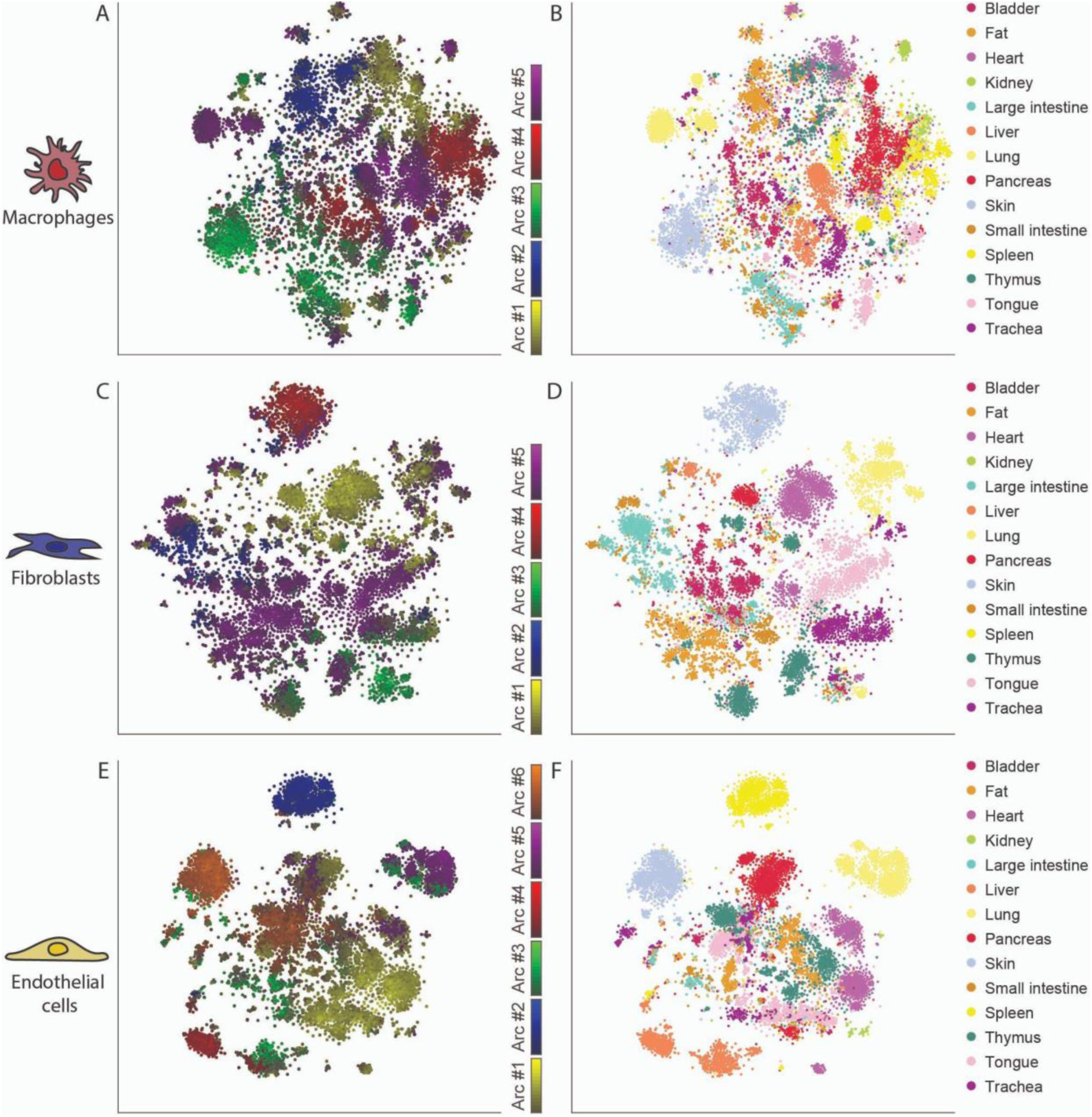
Tissue and archetype distance distribution in t-SNE space. Visualization of macrophage (A-B), fibroblast (C-D) and endothelial cell (E-F) gene expression in the low-dimensional t-SNE space representation. Cells are colored by their Euclidean distance to their corresponding archetype (left panels, brighter shade represents closer distance to archetype and grey represents generalist cells, see Methods: Analysis of tissue composition of archetypes) and by their tissue of origin (right panels).

**Figure S5:**
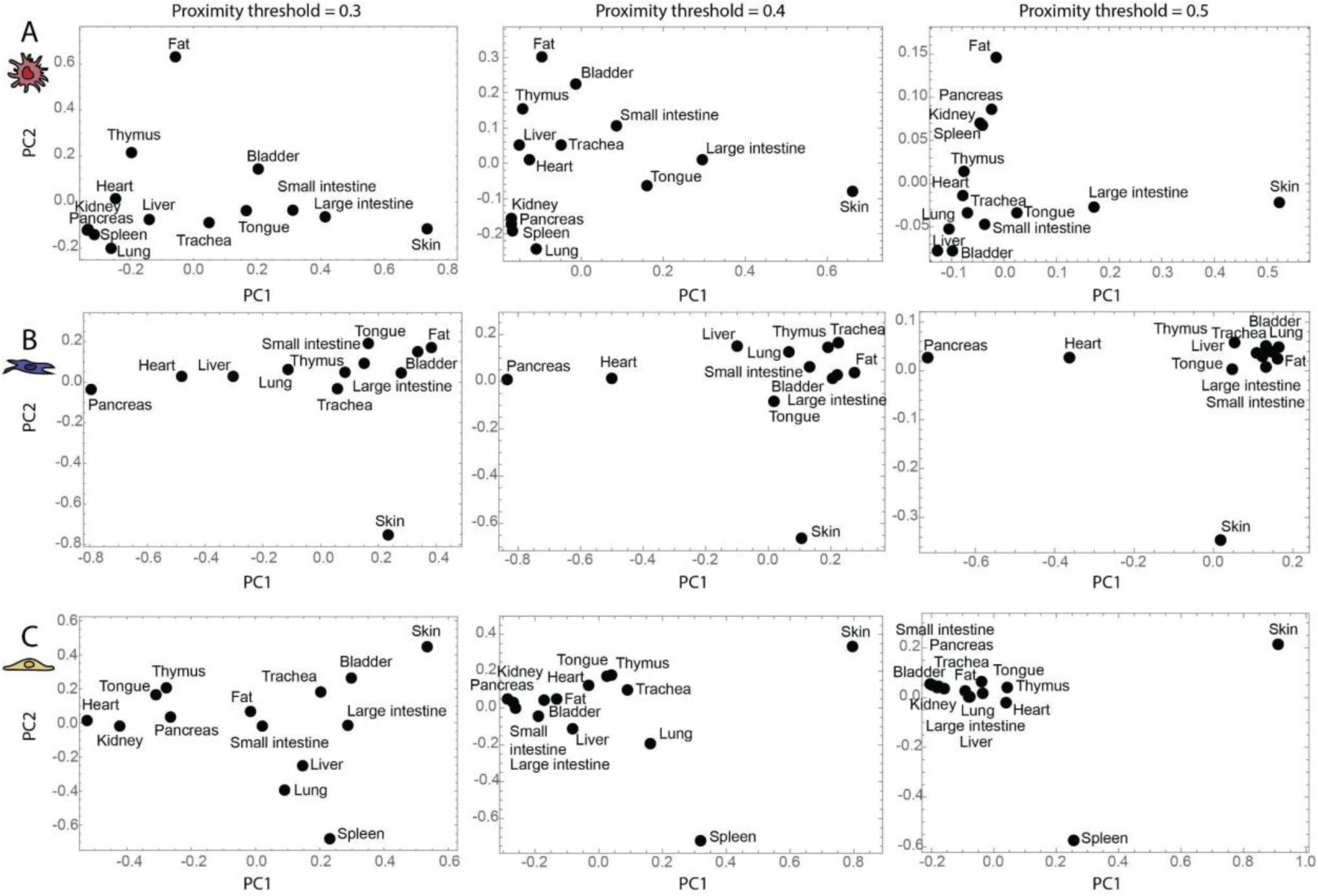
Proximity threshold sensitivity analysis for archetype distributions across tissues. PCA of archetype distributions across different proximity thresholds (0.3, 0.4, and 0.5). PCA was performed separately for each threshold on the tissue-level distribution of archetypes within macrophages (A), fibroblasts (B), and endothelial cells (C).

**Figure S6:**
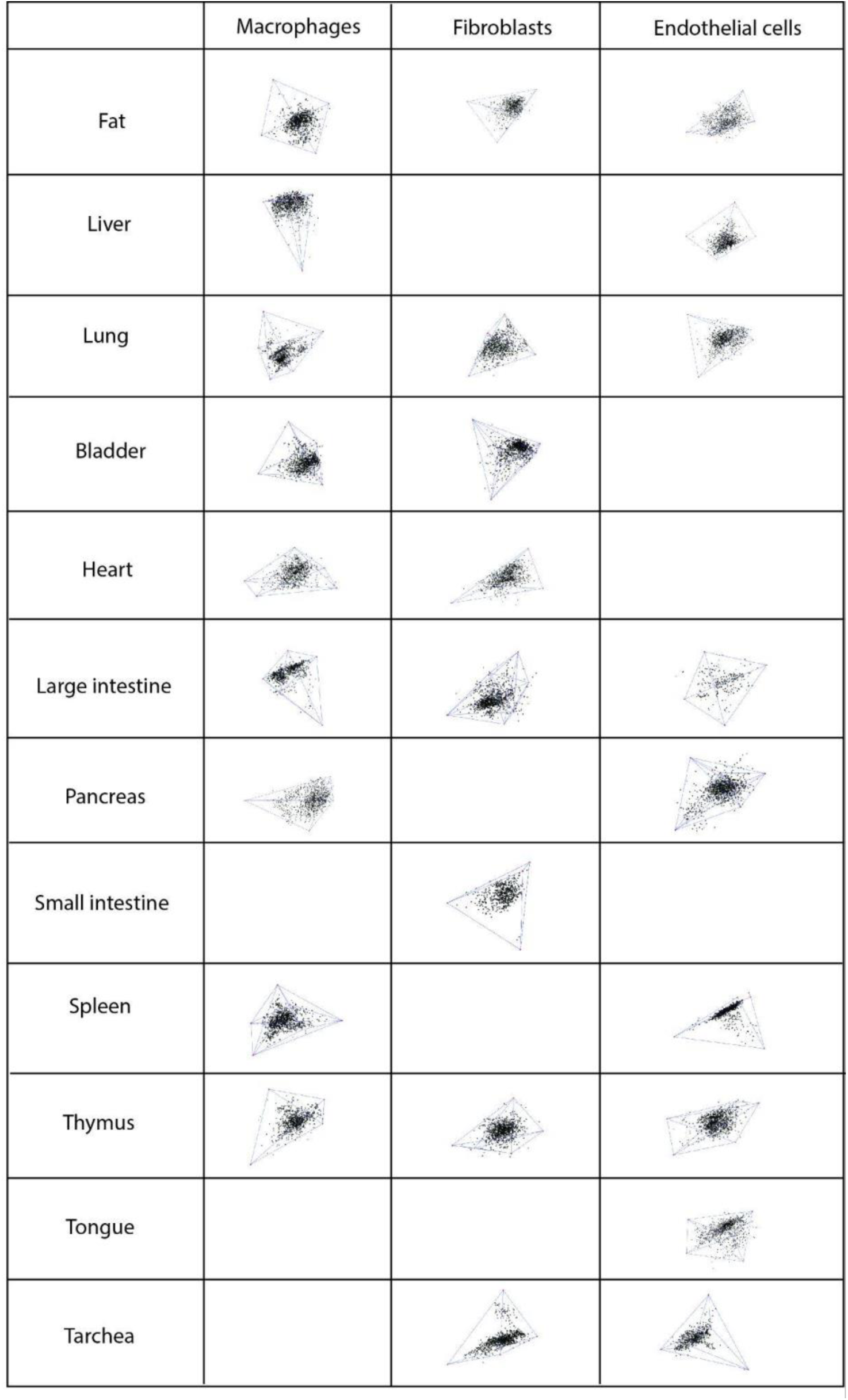
Archetype polytopes across tissues and cell populations in PCA space. Each cell in the table corresponds to a specific combination of tissue (rows) and cell population (columns: macrophages, fibroblasts, endothelial cells). Within each cell, black dots represent individual single cells and red dots indicate the archetype vertices defining the polytope, all visualized in PCA space. Empty cells indicate that no polytope was identified - either due to lack of statistical significance or absence of cells from that tissue in the analyzed dataset.

**Figure S7:**
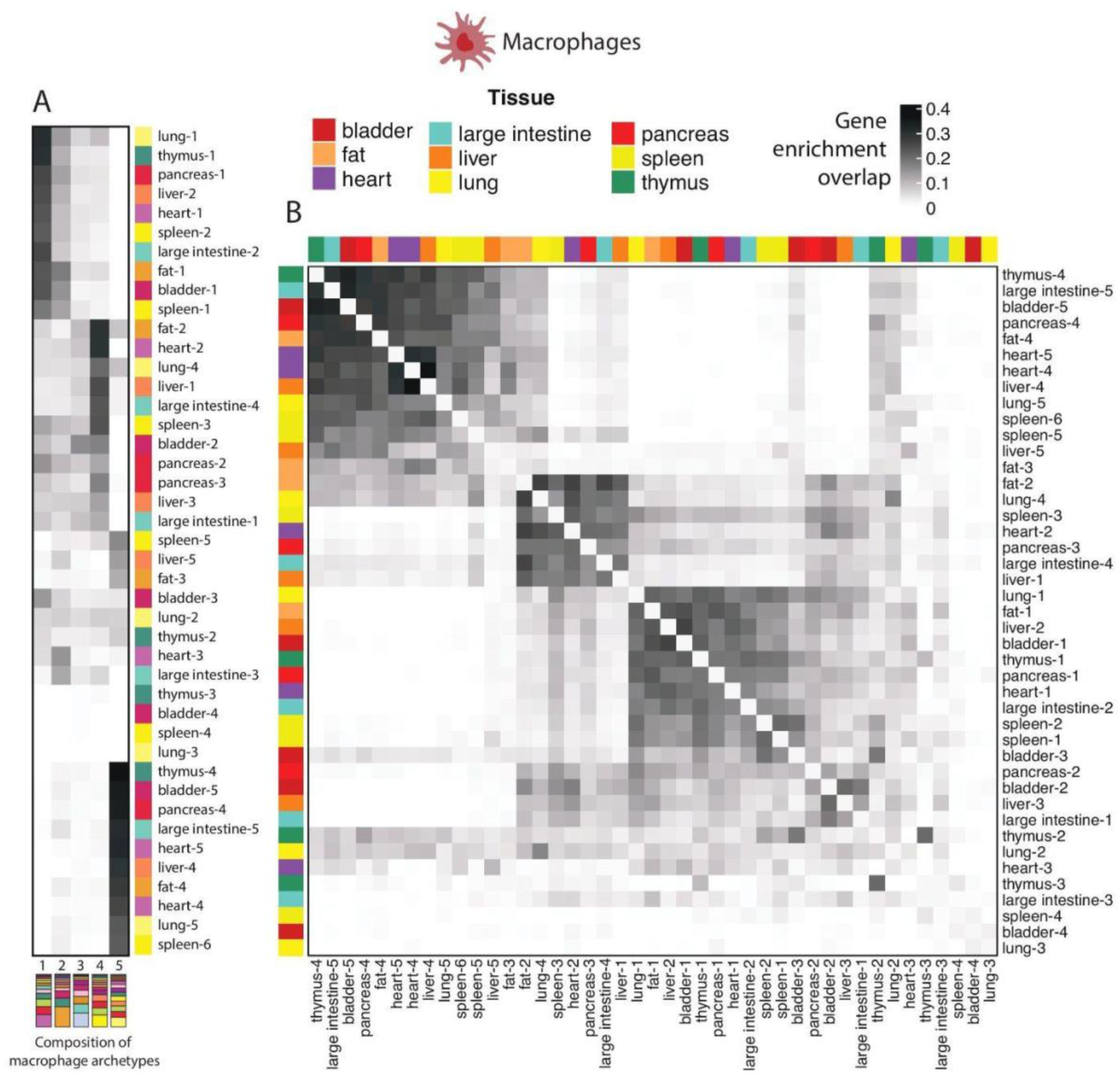
Archetype robustness and tissue-specific organization of macrophages. (A) Heatmap showing the overlap in enriched genes between individual-tissue macrophage archetypes (rows) and global macrophage archetypes (columns). Rows are clustered according to their similarity to the global archetypes. Darker shading indicates higher Jaccard overlap, with tissue origin for each archetype annotated by the color bar. (B) Heatmap comparing enriched-gene overlaps among all individual macrophage archetypes across tissues. Clustering is based on the similarity of enriched gene sets between archetypes. Tissue of origin is indicated by the color bars along both axes. This comparison highlights how macrophage archetypes from different tissues share or diverge in their gene enrichment profiles.

**Figure S8:**
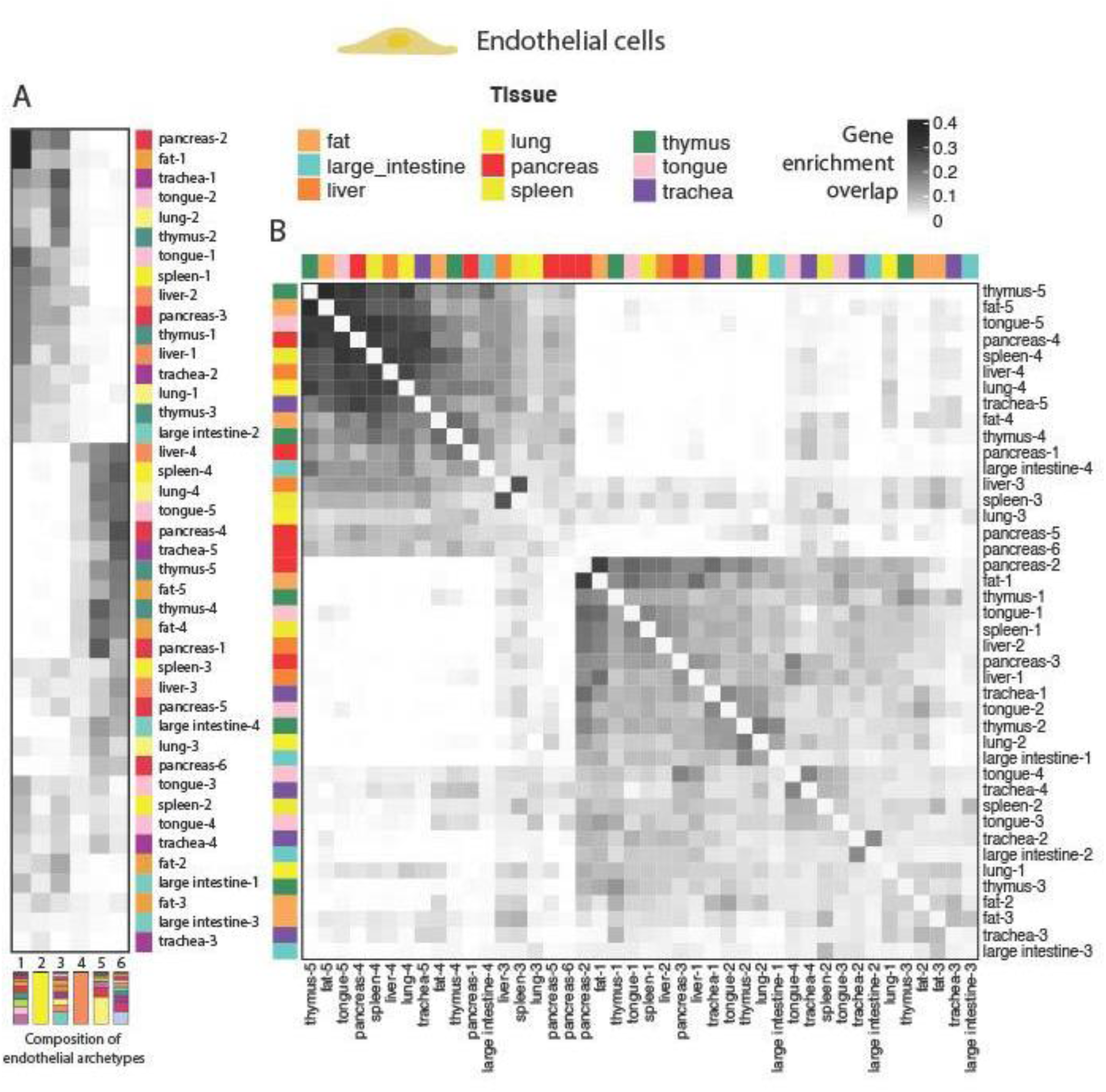
Archetype robustness and tissue-specific organization of endothelial cells. (A) Heatmap showing the overlap in enriched genes between individual-tissue endothelial cell archetypes (rows) and global endothelial archetypes (columns). Rows are clustered according to their similarity to the global archetypes. Darker shading indicates higher Jaccard overlap, with tissue origin for each archetype annotated by the color bar. (B) Heatmap comparing enriched-gene overlaps among all individual endothelial cell archetypes across tissues. Clustering is based on the similarity of enriched gene sets between archetypes. Tissue of origin is indicated by the color bars along both axes. This comparison highlights how endothelial archetypes from different tissues share or diverge in their gene enrichment profiles.

**Figure S9:**
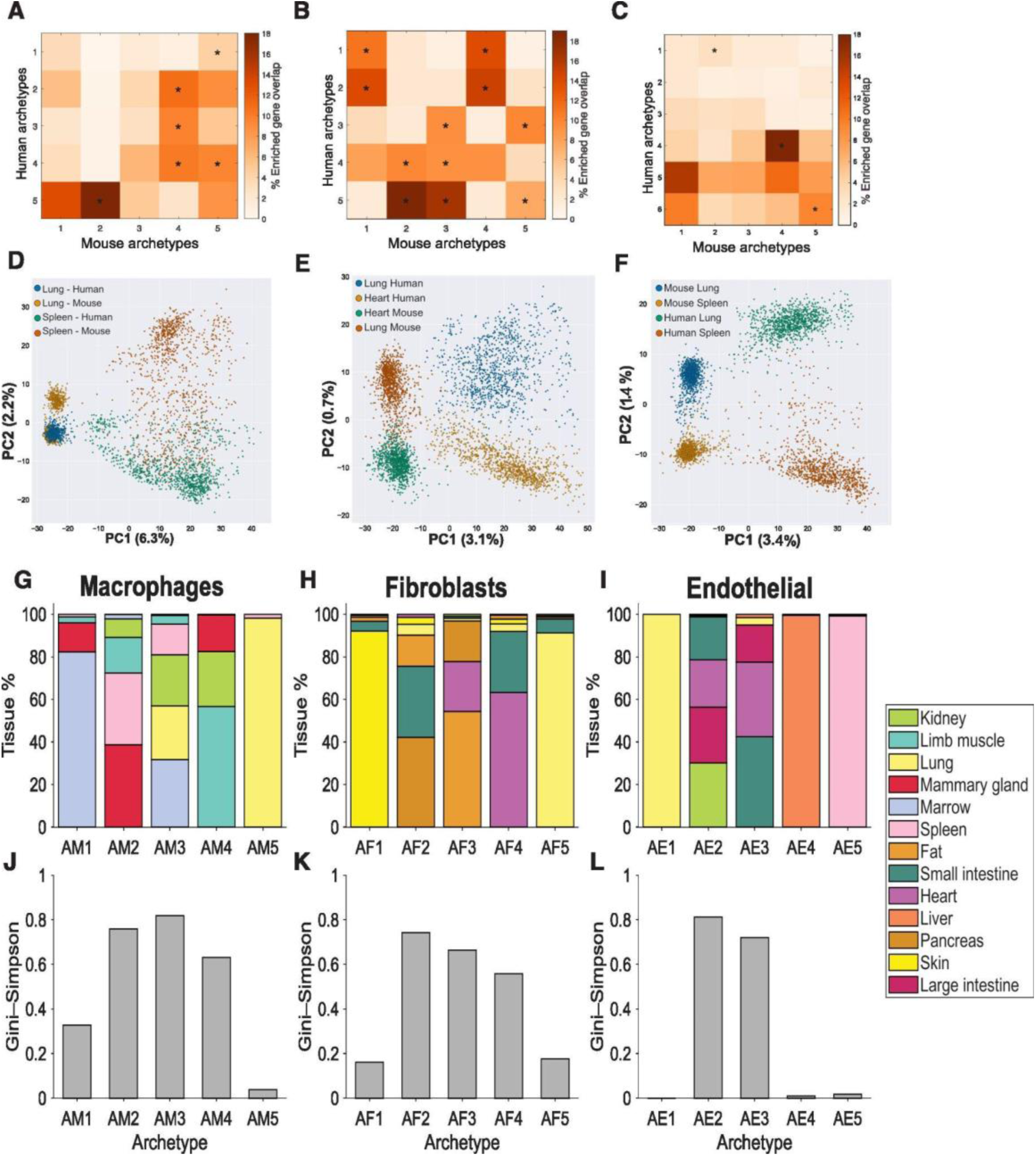
Cross-species comparison of supportive cell archetypes. (A-C) Heatmaps of gene enrichment overlap for macrophages (A), fibroblasts (B) and endothelial cells (C). Values show the percentage of enriched genes shared between each human archetype (rows) and mouse archetype (columns). Warmer colors indicate greater overlap. Asterisks mark mouse and human pairs with a statistically significant overlap (Methods). (D-F) Projection of mouse and human pairs of tissues onto the first two PCs in gene expression space for macrophages (D), fibroblasts (E) and endothelial cells (F). PCA was applied for gene expression of orthologues genes from the two tissues common to both species. (G-I) Tissue composition of mouse archetypes for macrophages (G), fibroblasts (H) and endothelial cells (I). Stacked bars display the proportion of cells from each tissue that contribute to every mouse archetype. (J-L) Gini-Simpson indices calculated from the tissue distributions for macrophage (J), fibroblast (K) and endothelial (L) archetypes.

